# A microwell platform for high-throughput longitudinal phenotyping and selective retrieval of organoids

**DOI:** 10.1101/2022.11.01.514733

**Authors:** Alexandra Sockell, Wing Wong, Scott Longwell, Thy Vu, Kasper Karlsson, Daniel Mokhtari, Julia Schaepe, Yuan-Hung Lo, Vincent Cornelius, Calvin Kuo, David Van Valen, Christina Curtis, Polly M. Fordyce

## Abstract

Organoids are powerful experimental models for studying the ontogeny and progression of diseases including cancer. Organoids are conventionally cultured in bulk using an extracellular matrix mimic. However, organoids in bulk culture physically overlap, making it impossible to track the growth of individual organoids over time in high throughput. Moreover, local spatial variations in bulk matrix properties make it difficult to assess whether observed phenotypic heterogeneity between organoids results from intrinsic cell differences or microenvironment variability. Here, we developed a microwell-based method that enables high-throughput quantification of image-based parameters for organoids grown from single cells, which can be retrieved from their microwells for sequencing and molecular profiling. Coupled with a deep-learning image processing pipeline, we characterized phenotypic traits including growth rates, cellular movement, and apical-basal polarity in two CRISPR-engineered human gastric organoid models, identifying genomic changes associated with increased growth rate and changes in accessibility and expression correlated with apical-basal polarity.

## Introduction

The advent of 3D organoid culture methods that recapitulate tissue structure, multi-lineage differentiation, pathology, and disease phenotypes while retaining the tractability of *in vitro* systems has revolutionized the study of various diseases, including cancer (*1–6*). Key strengths of organoids over conventional transformed 2D cell lines for studying cancer include the greater similarity of 3D versus 2D models to *in vivo* settings and the relatively clean genomic background of healthy primary organoids which can be engineered via CRISPR/Cas9 editing or other techniques to harbor alterations in oncogenes or tumor suppressors that promote malignant progression (*7–9*). Several studies have demonstrated the transformation of normal colon, stomach, and pancreas organoids into invasive carcinomas via the simultaneous introduction of multiple oncogenic ‘hits’ (*7, 8, 10–12*).These minimally transformed forward-genetic models have yielded insights into the requirements for transformation and their tissue-specific functional consequences.

For example, we recently developed individual and sequentially engineered organoid models to study the earliest mutational events during gastric tumorigenesis (*7, 13*). Gastric cancer is the third leading cause of cancer mortality worldwide and a major public health burden. Owing to limited screening modalities and late presentation of clinical symptoms, gastric cancer is commonly detected at an advanced stage, where treatment options are limited, emphasizing the need for earlier detection and robust biomarkers (*14*). We sought to recapitulate molecularly defined subgroups of disease via CRISPR/Cas9-mediated editing of genes commonly altered in human gastric cancers, including *TP53*, which is altered in 70% of chromosomally instable (CIN) gastric cancer (*15*), and *ARID1A*, which is altered in 50% of all cases and enriched in the microsatellite instable (MSI) and Epstein-Barr Virus (EBV) subgroups (*15, 16*). We first biallelically inactivated *TP53* via CRISPR/Cas9 in normal human gastric organoids and established clonally derived TP53 knockout (KO) lines. In these TP53-/- lines, we knocked out *ARID1A*, yielding TP53/ARID1A double knockout (DKO) lines that exhibited morphologic dysplasia, tumorigenicity, and mucinous differentiation, features that were not seen in TP53KO organoids (*7*). Additionally, through *in vitro* evolution of *TP53*-deficient gastric organoids, we demonstrated that this single initiating genetic insult is sufficient to recapitulate many molecular features of the CIN subgroup of gastric cancer while remaining morphologically similar to normal gastric organoids (*13*).

These single and double KO organoids thus represent powerful models of the earliest stages of human cancer, corresponding to pre-malignant and malignant states, respectively, and are ripe for further characterization. However, to date, the phenotypic characterization of single organoids has remained challenging as most techniques suitable for quantifying the growth of 2D cell cultures generally do not extend to 3D and many methods were designed for bulk populations (*17–20*). The most common strategy for culturing organoids in 3D involves resuspending dissociated cells in Matrigel or Cultrex Basement Membrane Extract (BME), commercially available extracellular matrix (ECM) mimics (*21*). Using this method, organoids are formed from clusters of aggregated cells rather than from a single cell that expands independently, making it difficult to determine whether an observed phenotypic trait reflects the stochasticity of small deposited cell populations or is intrinsic to individual cells at the start of organoid growth (*22, 23*). Distinguishing intrinsic from extrinsic heterogeneity is further complicated by variations in the organoid’s growth environment, as organoids seeded close to other organoids may be affected by cell-cell paracrine or juxtacrine signaling (*24, 25*).

Additionally, the position relative to the margins of the ECM hemisphere can impact the diffusion of growth factors and/or drugs (*26, 27*). Finally, the same observed bulk growth differences could result from differences in the median organoid growth rate (*28*) or the fraction of cells that grow, complicating the interpretation of such data.

Microfabricated microwells represent an alternative to bulk culture methods and have been used in the materials science and tissue engineering fields for 3D cell culture (*18, 22, 29–31*).However, most methods reported to date rely on cellular aggregation to generate spheroids (*22, 29, 31–35*) rather than growing organoids from a single cell, precluding investigations into underlying cellular and phenotypic heterogeneity. In addition, unlike in 2D cell lines, image analysis pipelines to quantitatively track the growth trajectories and other phenotypic traits of live 3D organoid cultures over time are less comprehensive (*31, 36, 37*). Finally, most platforms lack the ability to selectively retrieve organoids of interest for downstream investigation or require complex and expensive instrumentation, hindering efficient linkage of phenotype to genotype (*30*).

Here, we present an open-source microwell platform and image-processing pipeline that enables the characterization of a variety of phenotypes for thousands of single organoids in parallel under near-identical conditions. These microwells are easy to fabricate and integrate into traditional cell culture workflows and can be imaged using standard inverted microscopes. Applying this new microwell platform to two established CRISPR-engineered organoid models of gastric tumorigenesis, representing pre-malignant (TP53KO) and malignant (DKO) states, we quantify cell size and position over time to determine single organoid growth rates and migration patterns via time-lapse imaging and a neural network-based image analysis module (*38, 39*) for nearly 100,000 cell trajectories and over 8,000,000 microwell images. By fluorescently labeling nuclei and actin in both engineered organoid lines, we quantify the proportion of cells with abnormal apical-basal polarity, a hallmark of malignant transformation (*40, 41*), and demonstrate an enrichment in the DKO relative to the TP53KO model. We further implemented a mechanism to selectively retrieve specific organoids of interest for downstream characterization to facilitate efficient linkage of phenotype to genotype. With this, we investigated the molecular features associated with this phenotype by retrieving individual DKO organoids with normal versus disordered polarity and performing dual chromatin accessibility and transcriptomic profiling. These analyses implicate changes in accessibility amongst cell adhesion genes in apical-basal polarity. We anticipate that this microwell platform and associated analysis pipeline can readily be integrated within existing culturing protocols to interrogate the molecular and phenotypic features across a broad range of organoid and spheroid models.

## Results

### Organoid models of gastric tumorigenesis and growth rate measurements in bulk culture

Here, we use two established organoid models of gastric tumorigenesis based on single and combinatorial gene-editing via biallelic inactivation of the tumor suppressor, *TP53* (TP53KO) or dual inactivation of *TP53* and *ARID1A* (DKO) (*7, 13*) (**Fig. 1A**). The *TP53/ARID1A* DKO exhibited malignant phenotypes, including mucinous metaplasia and capacity for *in vivo* tumor growth upon xenotransplantation in mice, implying this line has undergone malignant transformation (*7*). In contrast, the TP53KO gastric organoids exhibit normal morphology and are not tumorigenic even after long-term *in vitro* evolution despite harboring a constellation of copy number variants associated with gastric cancer, implying this line mimics a pre-malignant state (*7, 13*). These defining features render these excellent models in which to perform the systematic and quantitative characterization of additional phenotypes within and between models, including growth rates and the prevalence of apical-basal polarity.

**Figure 1.**
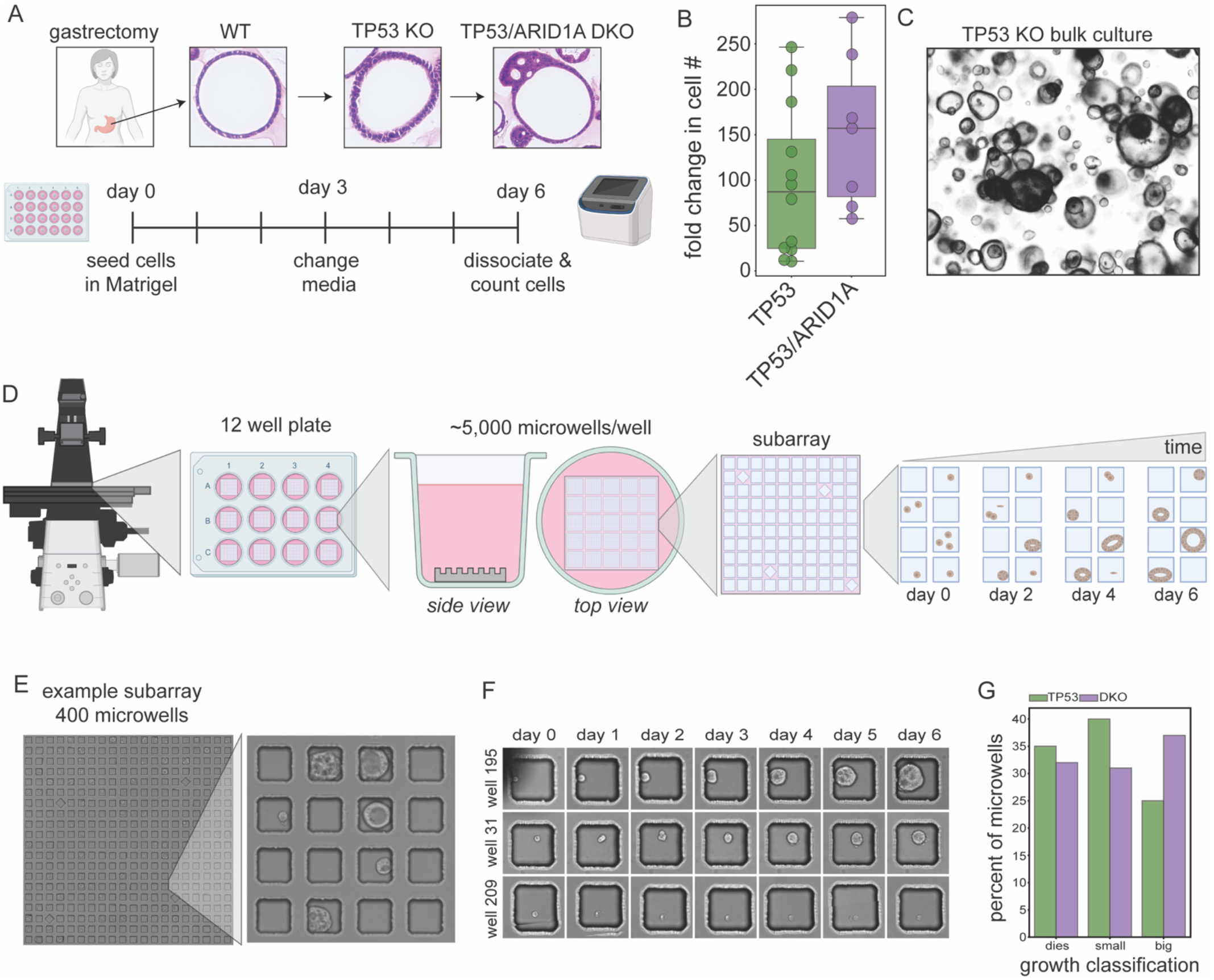
A microwell platform for high-throughput phenotyping of gastric cancer organoids. **(A)** Sequential genome editing of TP53 and ARID1A in human gastric organoids revealed progressive phenotypic changes. Loss of epithelial apical-basal polarity was associated with TP53/ARID1A DKO organoids. Organoid cultures were seeded in Cultrex BME, and allowed to grow for 6 days before bulk measurements of growth and microscopy examination. **(B)** DKO organoids appeared faster than TP53KO organoids in bulk measurement when comparing fold change in cell number from start to end of passage but the difference was not significant (Wilcoxon rank-sum test). **(C)** Example image of TP53 organoids grown in bulk culture, illustrating challenges with isolating individual organoids. **(D)** Schematic of the initial microwell experiments using brightfield microscopy. Each microwell array was placed onto a well of a 12-well tissue culture plate. Single cells dissociated from organoids were then seeded into the microwells and allowed to expand. **(E)** An example image of 100μm microwell subarrays with organoids at the end of a 6-day timecourse. **(F)** Growth trajectories of single cells seeded in microwells, demonstrating heterogeneity in growth patterns. **(G)** Manual classification of growth trajectories for single cells by organoid size at end of time course (small = <25% of well occupied, big = ≥25% of microwell occupied). A greater proportion of DKO organoids were classified as “big” compared to TP53KO.

As reported previously (*7*), images from bulk cultures grown in Cultrex BME revealed qualitative phenotypic differences between these gastric organoid lines, with increasing cellular disorganization for organoids lacking either *TP53* or both *TP53* and *ARID1A* (**Fig. 1A**). Changes in cell growth, assessed by seeding bulk cultures with the same number of cells and then quantifying the fold-change in the number of viable cells after 14 days of passaging in conditioned media, also suggested that DKO organoids were more proliferative than TP53KO organoids in bulk culture **(Fig. 1B**), consistent with previous observations (*7*). However, these bulk culture methods could not determine: (1) if fold-change increases resulted from individual cells growing at a faster rate or from differences in the fraction of cells that grow, (2) whether the culture contained subpopulations with different growth behaviors, or (3) whether any observed subpopulations resulted from intrinsic differences between cells or cellular microenvironment (*e.g*. proximity to other organoids or to media surrounding the Cultrex BME boundary). Finally, organoids in bulk culture physically overlap and can merge with one another **(Fig 1C**), making it difficult to track the growth of individual organoids or isolate individual organoids of interest for downstream analysis.

### Microwell arrays for high-throughput phenotyping of single organoids over time

To address these challenges, we developed a microwell platform to perform time-resolved phenotyping of thousands of organoids in parallel under near-identical conditions (**Fig. 1D**). Each single-layer PDMS microwell device contains arrays of 2,500-10,000 microwells (either 100 × 100 × 80μm or 200 × 200 × 80μm, length × width × depth) placed directly at the bottom of each well (“macrowell”) of a 12 well culture plate (**Fig. 1D; Supplementary Fig. S1**). Devices are fabricated by spin coating polydimethylsiloxane (PDMS) onto master molds, thereby ensuring uniform thickness and enhancing image quality (**Supplementary Fig. S2**). To facilitate unique microwell indexing during downstream image processing, microwells are grouped within subarrays of 20 × 20 (100 μm) or 10 × 10 (200 μm) microwells, with a pattern of rotated microwells that uniquely identifies each subarray (**Supplementary Fig. S3**). The 100 μm diameter microwells are optimized for high-throughput imaging of thousands of organoids in the same experiment, while 200 μm diameter microwells are best suited for retrieval of organoids with phenotypes of interest for single-organoid sequencing. All microwell devices were plasma-treated and coated with 0.5% BSA to render them hydrophilic.

Initial measurements of single-cell occupancy as a function of starting cell concentration for TP53KO cell lines established optimal concentrations of 6000 cells/mL for 100 μm microwells (26.15% of wells with a single cell) and 400 cells/mL for 200 μm microwells (31.33% of wells with a single cell), respectively; the fraction of microwells containing a given number of cells was well-fit by a Poisson distribution, consistent with expectations for stochastic loading (**Supplementary Fig. S4**).

For initial experiments, we seeded microwell arrays with single cells dissociated from TP53KO organoids and then imaged microwells daily via tiled bright field imaging (**Fig. 1E**). To visualize single organoid growth over time, we: (1) stitched tiled images into a single image per macrowell per timepoint, (2) rotated stitched images to position microwell array edges parallel with image edges, (3) manually determined corner locations for each macrowell, and then (4) extracted individual microwell images from the rotated arrays by their position relative to the corner locations (**Supplementary Fig. S3**). We then manually inspected a subset of 100 microwells for each organoid line. Organoids grown in microwells appeared phenotypically similar to their bulk culture counterparts, with circular and cystic structures (**Figs. 1E**). However, individual organoids grown from single cells under identical experimental conditions often showed dramatically different growth behavior: while some organoids showed rapid growth after seeding, defined as growth to ≥25% of the microwell area (25/100 TP53; 37/100 DKO) (**Fig. 1F**, top), others either grew very little (<25% of the microwell area; 40/100 TP53; 31/100 DKO) (**Fig. 1F**, middle) or showed signs of cell death/apoptosis (35/100 TP53; 32/100 DKO) (**Fig. 1F**, bottom). An approximately equal proportion of TP53KO and DKO organoids exhibited growth; based on our manual classification, a greater proportion of DKO organoids showed larger and more rapid growth (≥25% of the microwell area) compared to TP53 (37% versus 25% of measured organoids, respectively) (**Fig. 1G**).

### High-throughput fluorescence imaging combined with deep learning tracks thousands of cells in parallel

Brightfield microscopy made it possible to qualitatively assess organoid growth, but manual classification methods were both time-consuming and confounded by a high degree of subjectivity. In addition, brightfield images lacked sufficient resolution to accurately determine whether observed growth was a result of cell division or simply an increase in organoid volume due to osmotic swelling of the lumen and the cells (*42, 43*). To distinguish between these possibilities and simultaneously gain information about nuclear localization within organoids, we engineered both TP53KO and DKO gastric organoid lines to express a nuclear fluorescent reporter by lentivirally inserting a mCherry-tagged copy of histone 2B (mCherry-H2B) (**Fig. 2A**); in parallel, we profiled the distribution of cytoskeletal proteins using live-cell labeling of actin or tubulin (*e.g*. SiR Actin Kit from Cytoskeleton Inc.). To determine optimal conditions for fluorescence imaging and pilot new analysis pipelines, we loaded TP53KO and DKO engineered cells (at a concentration of 4000 cells/mL) into 100 μm microwells across three separate replicate experiments. The first and second experiments took place after 5 and 7 months of continuous passaging in conditioned media; the third experiment took place after ~8 months of continuous passaging followed by a freeze-thaw cycle (required due to COVID pandemic-related shutdowns) and an additional ~1 month of passaging post freeze-thaw (**Fig. 2A**). After loading, we mounted the entire plate assembly on an automated microscope with an incubation chamber and collected tiled images across the device in the bright field, mCherry, and Cy5 (for fluorophore-tagged actin molecules) channels at 2-hour intervals over 5 days for each experiment (**Fig. 2A**). We then performed stitching, rotation, and microwell extraction for each imaging channel, similarly to the initial brightfield tests (ref. Methods). Time-course image processing for each well yielded 8,014,800 microwell images.

**Figure 2.**
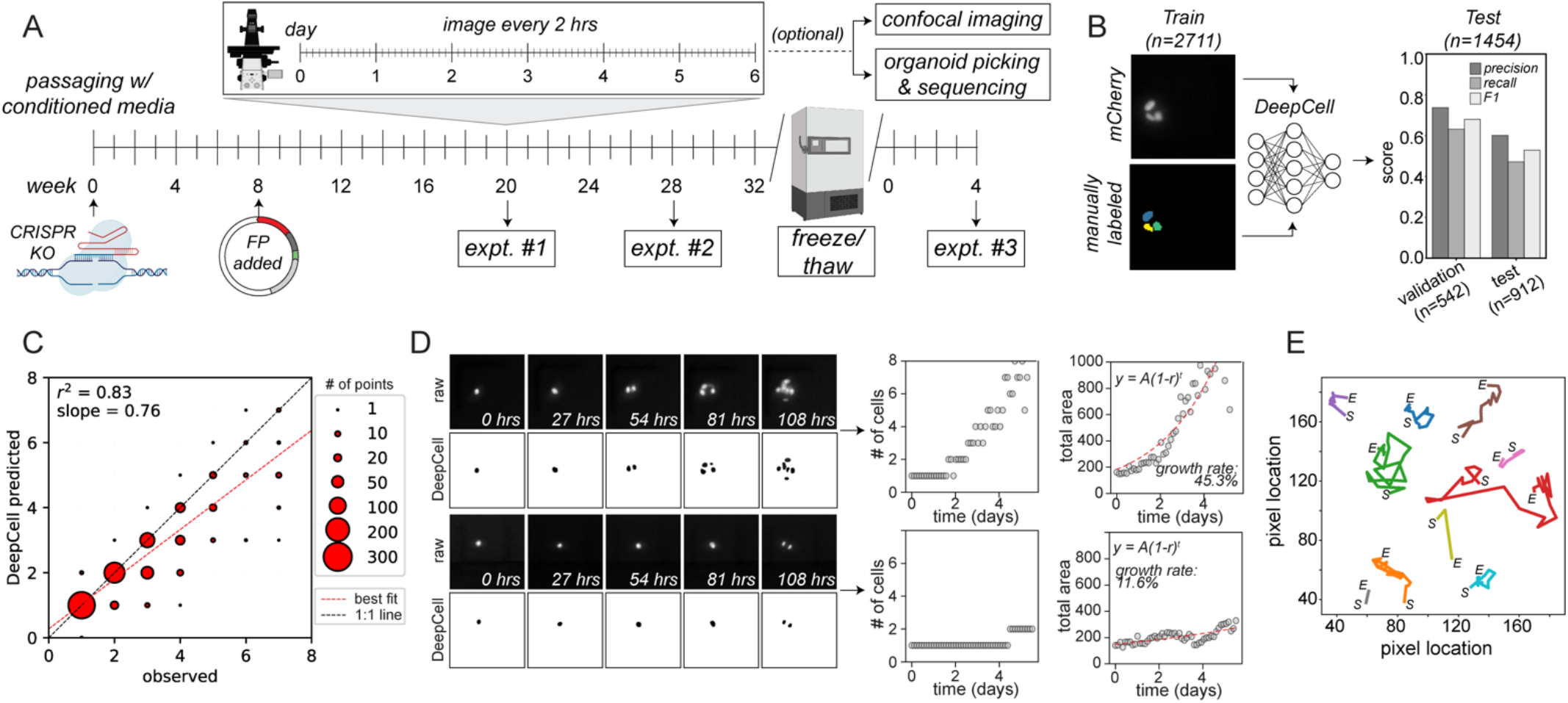
Neural net accurately identifies cells within fluorescence images to enable high-throughput image processing and organoid growth quantification. **(A)** Experimental timeline for cell engineering and imaging experiments. **(B)** Pipeline for model training and validation (left) and performance metrics (right). **(C)** Scatter plot comparing the number of model-predicted cells and manually annotated cells per microwell (N = 912 total microwells). Marker size indicates the number of events, dashed red line indicates linear regression, and dashed black line indicates 1:1 line. **(D)** Example raw and model-predicted images (left) and model-annotated cell numbers and areas over time (right) for 2 microwells over 108 hours of imaging (left). Red line shows the exponential growth equation using the indicated best fit parameters; growth rate is reported based on the % daily change in area). **(E)** Example single cell position traces over time.

To efficiently extract information about the number, size, and relative positions of cells within this large imaging dataset, we turned to a neural network originally designed for optical microscopy-based image segmentation of cells grown in 2D monolayers (*38, 39*). To test whether and under what conditions this deep learning model can recognize and track cells within 3D organoids, we manually labeled individual mCherry-tagged nuclei positions in 2711 images from the first experiment, used 2169 of these images and a transfer learning approach to train an organoid-aware version of the model, and validated performance with the remaining 542 images (Methods). To evaluate generalizability across experiments, we quantified model performance based on the number of accurately identified cells within an additional 912 manually labeled ‘test’ images taken from all 3 experiments (306, 348 and 258 images from experiments #1-3 respectively) (**Fig. 2B**). The deep learning model counts showed strong concordance with manual counts within the entire 5-day duration of the experiments (*R^2^* = 0.83 for the three combined experiments; **Fig. 2C; Supplementary Fig S5A** for per-experiment correlation). Precision and F1 scores were generally higher than recall for both validation and test sets, with the validation set displaying higher scores across performance metrics, as expected (**Fig 2B; Supplementary Fig. S5B**). Examination of error type revealed substantially more merges (in which 2 labeled cells were predicted by the model to be a single cell) compared to splits (where a single labeled cell was predicted to be 2 cells).

Further visual inspection of images showing regions identified as cells by the deep learning model confirmed the accuracy of automated annotations over time (**Fig. 2D**). Identifying all cells from microwells within a single macrowell required an average of 145 seconds for model processing per time point. Fitting the model-annotated cell area and cell number over time for each microwell to an exponential growth curve yielded quantitative growth rate estimates (**Fig. 2D**); linearly interpolating the centroid position of single cells over time returned a lower bound for the distance traveled by a single cell prior to the first cell division (**Fig. 2E**).

### P53/ARID1A DKO cells grow at faster rates and higher occupancies decrease growth rates

Using the trained deep learning pipeline for 3D cultures, we then quantified organoid growth rates across all three initial fluorescence microscopy experiments (**Fig. 3A**). In total, the three experiments profiled growth rates for 5812, 1328, and 3679 loaded microwells in experiments #1-3, respectively, with the number of cells loaded per microwell following stochastic Poisson distribution (**Supplementary Fig. S6A**). Somewhat surprisingly, per-organoid growth rates for both organoid lines decreased as the number of cells initially seeded within the microwell increased, suggesting that increased paracrine or juxtacrine signaling between cells in close proximity does not enhance growth (**Fig. 3B**). We restricted analysis downstream to microwells seeded with a single cell. Similar to growth rates observed in bulk cultures, the single-organoid growth rates of the DKO line were higher than those of the TP53KO line (**Fig. 3C**). However, we observed ~2-fold variability in growth rate across experiments, presumably due to differences in the composition of conditioned media across batches (*44*). Despite this, DKO organoids consistently grew faster than TP53KO organoids within the same experiment (**Fig. 3C**), establishing that the higher fold-change in cell numbers for DKO organoids in bulk experiments (**Fig. 1B**) is due to an enhanced growth rate in individual organoids rather than growth of a larger proportion of cells. The single-organoid growth rates for the same organoid line across different macrowells within the 12 well imaging plate varied only slightly, confirming lack of macrowell-specific growth effects (**Fig. 3D**).

**Figure 3.**
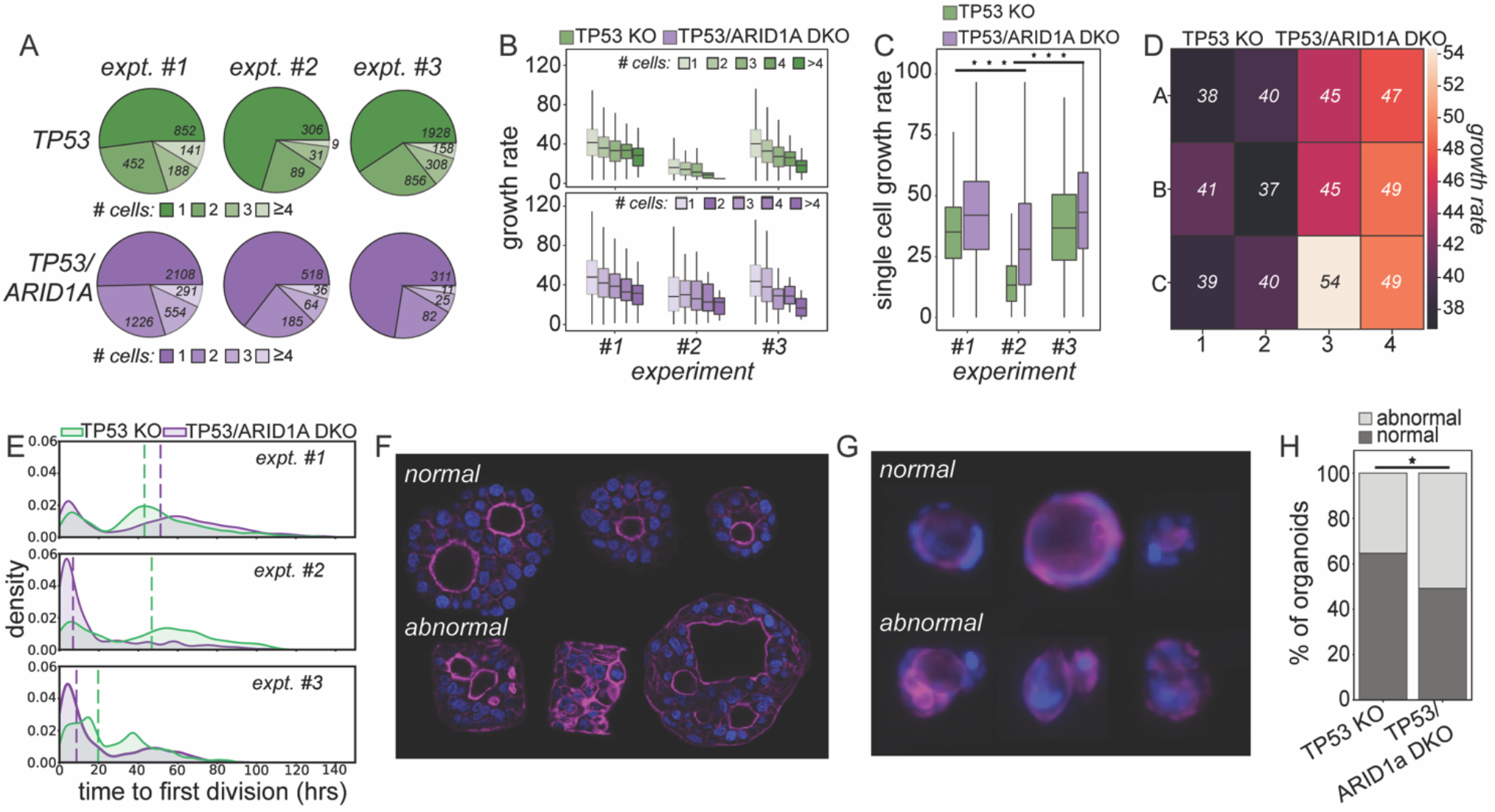
Microwell experiments and analysis reveal differences in growth rates and epithelial apical/basal polarization. **(A)** Pie charts indicating the number of TP53KO and TP53/ARID1A DKO cells loaded per microwell that exhibit growth across each experiment. **(B)** Growth rates as a function of the number of cells per microwell across each experiment for TP53KO (top, green) and TP53/ARID1A DKO (bottom, purple) organoid lines. **(C)** Growth rates for all single cells across each experiment for TP53KO (green) and TP53/ARID1A DKO (purple) organoid lines. Wilcoxon rank-sum tests yielded significant differences between experiment #2 and either experiment #1 or #3 for both within-TP53KO and within-DKO comparisons (*** denotes Bonferroni-adjusted p-value <0.001). **(D)** Heat map showing median growth rates (% daily change in cell area) for all microwells within macrowell A1 for experiment #1; median growth rates are indicated within each well. **(E)** Kernel density plots showing the time to first cell division for microwells containing a single cell from TP53KO or TP53/ARID1A DKO organoid lines; dashed lines indicate median time to first cell division. **(F)** Example confocal images of normal and abnormal polarity organoids. **(G)** Example fluorescent images of normal and abnormal polarity organoids. **(H)** Percentage of organoids with normal vs. abnormal polarity TP53KO vs. DKO lines. Fisher’s exact test yielded a significant difference between the two groups (p-value=0.03).

To calculate the time required for cells to complete a first division, we also determined the time point at which 2 cells were first identified within microwells originally seeded with a single cell. In all 3 experiments for both P53KO and DKO mutants, we observed a bimodal distribution in which some cells divided soon after initial seeding (0-24 hours post-seeding) and the remainder took longer to divide (24-120 hours post seeding) (**Fig. 3E**). We also observed experiment to experiment variation in the time to first division. For example, most P53KO cells in experiment #2 began dividing soon after seeding, and only a small subset of cells divided at later time points, compared to experiment #1 in which the population was much more evenly distributed. The total distance moved exhibited a similar modal pattern to the time to first cell division, as cells that took longer to divide had more time for movement (**Supplementary Fig. S6B**). Total distance moved was moderately positively correlated with the time to first division (R^2^ = 0.3209; **Supplementary Fig. S6C**), and weakly negatively correlated with growth rate (R^2^ = 0.0148; **Supplementary Fig. S6D**). Cells tended to move the furthest immediately after seeding, and then settled down with decreasing movement before dividing (**Supplementary Fig. S6E**).

### Microwell enabled examination of single organoids with disordered polarity

Disordered apical-basal polarity is commonly seen in diverse epithelial cancers (*41, 45*) and is believed to be an early hallmark of gastric cancer (*45, 46*). Intriguingly, in previous work, we noted that TP53/ARID1A DKO gastric organoids exhibit a disordered polarity (*7*). To assess whether apical-basal polarity could be visualized in organoids grown in microwells, we performed confocal imaging of both TP53KO and DKO, which confirmed that both TP53KO and DKO lines contained organoids with disordered polarity. To systematically quantify changes in membrane organization, we visually inspected the final timepoint fluorescence images for 257 and 236 TP53KO and TP53/ARID1A DKO cells, respectively, from the first experiment and classified organoids as having either ‘normal’ or ‘abnormal’ apicobasal polarity. Classification of ‘abnormal’ polarity was based on the requirement that organoids display two of the following three characteristics: (1) disordered actin signal not restricted to organoid lumens, (2) lack of a central lumen ringed with actin, and/or (3) presence of multiple small, disorganized lumens (**Fig. 3F-G**). A greater proportion of the DKO organoids exhibited an ‘abnormal’ phenotype (50.8% vs. 35.4% in the TP53KO organoids, p = 0.03, Fisher’s exact test), suggesting that more complex engineered genotypes drive greater cellular disorganization (**Fig. 3H**).

### Chromosome alterations associated with enhanced organoid growth

The high-resolution phenotyping data from these initial three experiments (#1 - #3) provided evidence that subtle differences in experimental conditions (*i.e*. different media batches) between experiments can drive phenotypic differences large enough to obscure meaningful intrinsic biological variation (**Fig. 3C**). To systematically characterize variability within and across TP53KO versus DKO organoids, we performed three additional experiments under more tightly-controlled conditions (ref. Methods) in which all cells were passaged using a standardized chemically defined media (**Fig. 4A**); these experiments used arrays of 200 μm microwells to enable retrieval of organoids of interest using tubing with an outer diameter of 250 μm. In total, these experiments profiled 422, 147, and 1009 loaded microwell with growth in experiments #4-6, respectively (**Fig. 4B**).

**Figure 4.**
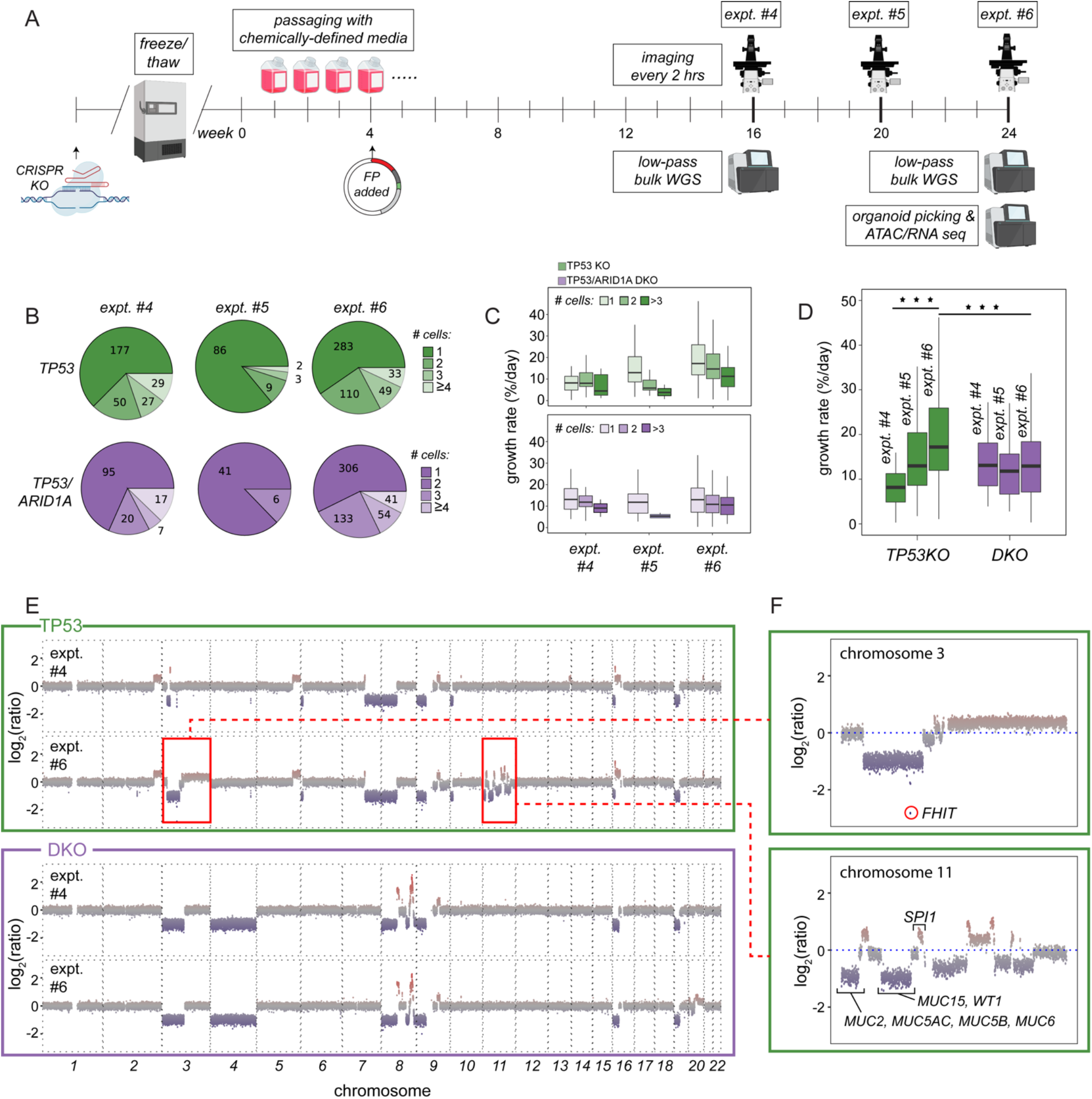
Microwell experiments under controlled experimental conditions reveal genomic changes associated with increased growth rates for TP53KO organoids. **(A)** Schematic of experiments #4 - #6 using chemically-defined growth media. TP53KO and DKO organoid lines were revived from cryopreservation and lentivirally transduced with H2B-mCherry at a defined MOI of 0.1 in 200μm microwells; shallow whole-genome sequencing was performed on bulk cultures at the start of experiments #4 and #6. (B) Pie charts indicating the number of TP53KO and TP53/ARID1A DKO cells loaded per microwell across each experiment. (C) Growth rates as a function of the number of cells per microwell across each experiment for TP53KO (top, green) and TP53/ARID1A DKO (bottom, purple) organoid lines. (D) Growth rates for all single cells across each experiment for TP53KO (green) and TP53/ARID1A DKO (purple) organoid lines. TP53KO organoids grew increasingly faster with each experiment while the growth rates of DKO organoids remained relatively constant (*** denotes Bonferroni-adjusted p-value < 0.001; Wilcoxon ranked-sum tests). (E) Whole-genome copy number plot for TP53KO (green box) and DKO organoids (purple box) from experiments #4 and #6. (F) Zoomed-in view showing copy number variations across chromosomes 3 and 11 for TP53 organoids at the start of experiment #6.

Similar to the previous experiments (**Fig. 3**), growth rates generally decreased as per-microwell occupancy increased for both TP53KO and DKO organoids (**Fig. 4C**). Most other image-based analyses such as distance moved and time to first division also recapitulated the observations made in **Fig.3** (**Supplementary Fig. S7**). The DKO organoid growth rates remained relatively constant across experiments (**Fig. 4D**), highlighting the consistency of the growth media composition. However, while TP53KO organoid growth rates were initially slower than those of DKO organoids, consistent with prior experiments, TP53KO growth rates increased steadily across experiments (p=3.48E-7, 2-tailed t test) until they eventually exceeded the DKO organoid growth rates by a significant margin (p = 0.025, 2-tailed t test) (**Fig. 4D**).

Previously, we observed that TP53KO organoids rapidly and continuously accumulate copy number variation (CNVs) and aneuploidy (*13*). To evaluate whether increased growth rates might be attributable to additional CNVs, we performed shallow whole-genome sequencing (sWGS) of both TP53KO and DKO organoid cells frozen during seeding of experiments #4 and #6. The sWGS for the TP53KO organoids revealed additional large-scale copy number alterations on chromosome 3 in experiment #6, as well as apparent chromosome shattering encompassing all of chromosome 11, that were not present in experiment #4 (**Fig. 4E**). By contrast, DKO cells showed no significant CNV changes across experiments (**Fig. 4E**), corroborating the previous observation that DKO organoids did not exhibit molecular signatures associated with the CIN subtype of gastric cancer, but instead displayed transcriptional profiles characteristic of the MSI subtype (*7*).

Next, we investigated regions affected by CNVs in TP53KO cells to identify specific genetic changes that could explain the observed increase in growth rate. While the TP53KO organoids from both experiment #4 and experiment #6 (passage #9 and #13 post-thaw) harbored focal deletion of FHIT locus, a common early alteration in this model and in gastric cancers (*13*), the entire chromosome 3p arm was deleted in experiment #6 (**Fig. 4F**). FHIT is a secondary regulator of DNA damage response commonly altered in TP53-null organoids (*13*), and in gastric cancer (*47, 48*), and the data suggests that the loss of both FHIT and TP53 likely drives rapid accumulation of additional CNVs. On chromosome 11, we observed loss of several regions containing multiple mucin genes (MUC2, MUC5AC, MUC5B, MUC6, and MUC15) (**Fig. 4F**) that are often dysregulated during malignant progression (*49–51*). For instance, abrogation of *MUC5AC* expression has been associated with increased cell proliferation and vascular invasion in gastric tumor cells (*52, 53*). In addition, we observed amplification of *SPI1* which is a proto-oncogene upregulated in various cancers and implicated in proliferation (*54–56*) (**Fig. 4F**). These alterations likely explain the increased growth rate observed in the TP53KO organoids over time.

### Sequencing of abnormal versus normal polarity organoids reveals changes in chromatin accessibility associated with morphology and adhesion

To explore the molecular changes driving differences in cell polarity, we extracted DKO organoids of interest from the microwell arrays for downstream sequencing using a syringe pump and 3D-printed microscope adapter (Methods; **Fig. 5A**). After staining organoids in microwells at the end of experiment #6 with a live-cell actin dye (ref. Methods), we retrieved ten organoids with normal polarity and ten with abnormal polarity (determined using the criteria described in Methods) from the DKO line (**Fig. 5B; Supplementary Fig. S8**). We then prepared libraries for single-organoid dual ATAC-seq and RNA-seq to identify differentially accessible regions (DARs) or differentially expressed genes (DEGs), respectively, that might explain differences in polarity.

**Figure 5.**
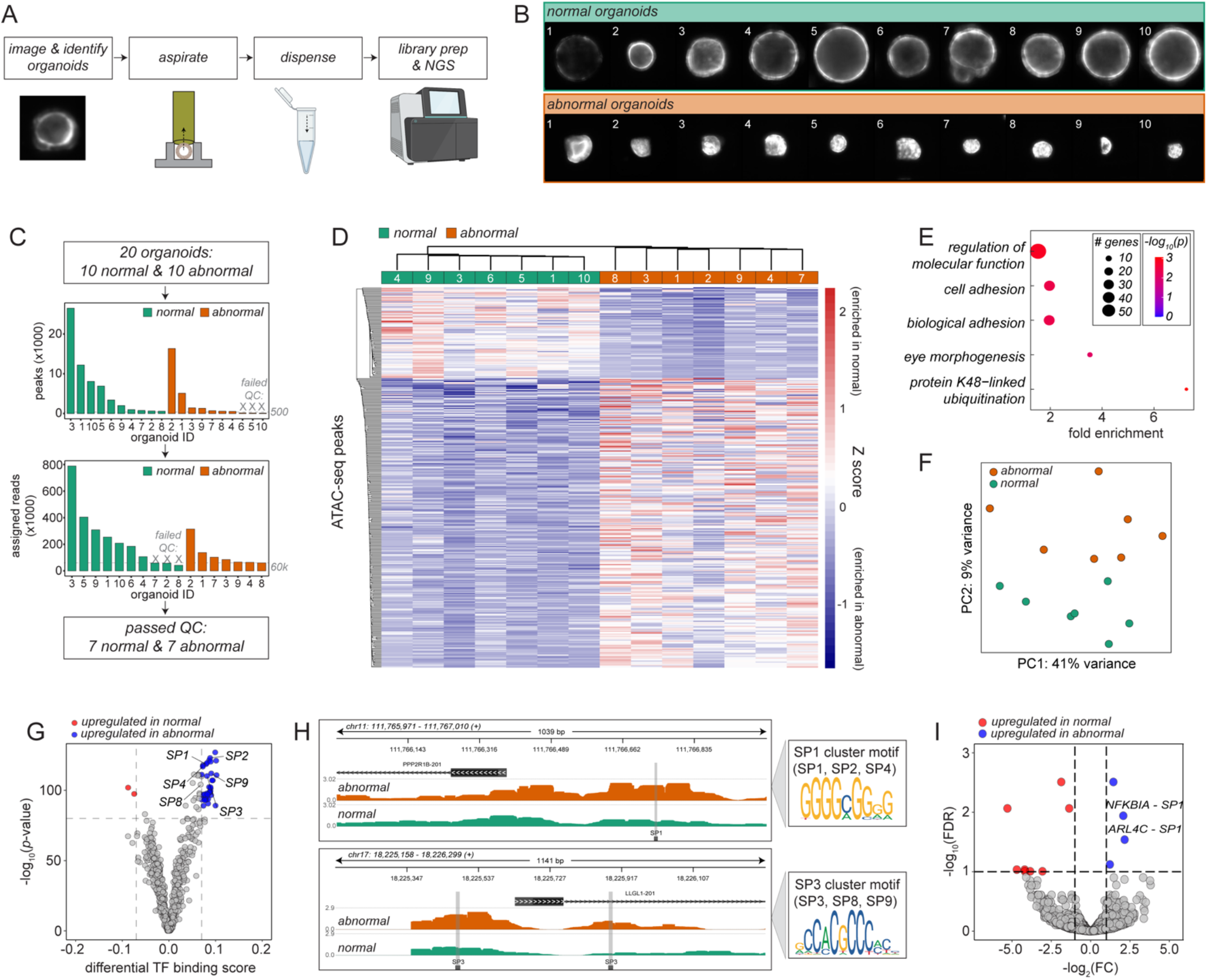
Single organoid retrieval and sequencing reveals changes in chromatin accessibility associated with changes in apical/basal polarity. **(A)** Schematic of experimental pipeline for single organoid retrieval. Single organoids were retrieved using a syringe pump connected to a 3D-printed microscope adapter via PEEK tubing and dispensed into a clean vial prior to preparing ATAC-seq and mRNA-seq libraries. **(B)** Images of all retrieved organoids. Top: organoids with normal polarity (*i.e*. a robust actin ring around the lumen). Bottom: organoids with abnormal polarity (*i.e*. disorganized actin signals). **(C)** ATAC-seq quality control pipeline eliminating samples with insufficient read depths or peaks. **(D)**Heatmap showing differentially-accessible peaks between normal and abnormal organoids. **(E)** Gene ontology analysis of differentially-accessible peaks showing enrichment of the peaks in pathways associated with cellular adhesion and morphology. **(F)** Principal component analysis of ATAC-seq data showing clear separation between normal and abnormal organoids. **(G)** Transcription factor footprinting analysis of ATAC-seq data showing enriched binding of SP-family transcription factors in abnormal organoids. **(H)** Representative plots showing increased binding of SP-family transcription factors in PPP2R1B and LLGL2 regulatory regions involved in regulating apical-basal polarity in epithelial cells. **(I)** Volcano plot showing differentially expressed genes in RNAseq data. Two of the four upregulated genes in abnormal organoids are known to harbor binding sites for SP-family transcription factors.

Given the limiting biomass derived from single organoids, we used a dual library preparation protocol developed for low-input specimens from Li *et al* (2021)(*57*). As the data quality was generally higher for ATAC-seq libraries compared to RNA-seq libraries, we focused on the ATAC-seq data initially. Following read deduplication and exclusion of six organoid samples with <60,000 uniquely mapped reads or <500 accessible ATAC-seq peaks, we profiled seven normal and seven abnormal polarity DKO gastric organoids with an average of 191,000 unique ATAC-seq reads and 4,425 accessible peaks per sample (**Fig. 5C**). Clustering based on ATAC-seq DARs clearly separated organoids with normal vs. abnormal polarity (**Fig. 5D**), and the DARs were enriched for gene ontology (GO) terms related to cell adhesion and morphogenesis (**Fig. 5E**). Principle component analysis (PCA) also revealed a clear separation between normal and abnormal polarity organoids (**Fig. 5F**). At a molecular level, transcription factor (TF) footprinting analysis of the ATAC-seq data indicated that organoids with abnormal polarity had increased accessibility in chromosomal regions bound by the SP family TFs (**Fig. 5G**), which are involved in maintaining epithelial cell polarity (*58, 59*) and are dysregulated in a variety of cancer types (*60, 61*). Consistent with this potential role, SP family TFs were more highly bound to upstream promoter regions of several genes such as *PPP2R1B* and *LLGL1* involved in establishing epithelial apical-basal polarity (**Fig. 5H**) (*59, 62, 63*). The RNA-seq data corroborated these findings, as multiple upregulated genes with known SP family target sites were differentially expressed in normal versus abnormal polarity organoids (**Fig. 5I; Supplementary Fig. S9**). These data thus suggest a plausible molecular basis for abnormal polarity in TP53/ARID1A DKO gastric organoids and demonstrate the feasibility and utility of molecularly characterizing single organoids with distinct imaging-based phenotypes.

## Discussion

Here, we describe an easy-to-use, open-source microwell platform for high-throughput, image-based phenotyping of organoids with the ability to retrieve single organoids of interest for additional downstream molecular profiling (*e.g*. single-organoid ATAC-seq or RNA-seq). As a first demonstration of this platform, we used an organoid-optimized deep learning model to identify and track nearly 100,000 individual cell trajectories from two engineering human gastric organoid model lineages within >8,000,000 microwell images. Using these data, we then quantified organoid growth rates and positions over time with high accuracy and granularity. These measurements led to the identification of specific molecular features associated with increased organoid growth and loss of apical-basal polarity.

This microwell platform offers multiple advantages over other existing organoid culture methods. In comparison to bulk culture, the microwell method combined with our custom open-source 3D image analysis module allows us to precisely measure quantitative phenotypes for individual organoids, such as single organoid growth rates, division times, and migration distances, rather than being restricted to bulk averages, such as fold change in cell number for an entire population (*64*). Unlike commercially available methods or closed microfluidic systems (*18, 65*), our microwell arrays do not require specialized instrumentation and are thus easy to adapt to existing cell culture workflows (*31*). In addition, we were able to generate complete organoids from single cells seeded in microwells (*32–35*), a particularly relevant advance for studying the early stages of tumorigenesis, in which a single transformed cell develops into a tumor. We also demonstrated that our microwell platform has comparatively higher throughput than other existing methods, being able to profile thousands of organoids in parallel as opposed to hundreds at a time (*29, 34, 37*). Finally, our microwell platform also facilitates retrieval of individual organoids of interest, which can then be either directly sequenced to investigate the molecular changes driving particular phenotypes or clonally expanded *in vitro* to generate sufficient biomass for additional phenotypic or molecular profiling.

In future work, we anticipate that the ability to profile single organoids will enable personalized drug screening in patient-derived tumor organoids. The microwell platform described here can be used to characterize the drug response and associated phenotypic changes of each organoid seeded in the microwells independently, offering the potential to screen patient-derived tumor organoids against a wide variety of drugs using much lower input materials. Additionally, the organoid retrieval mechanism described here confers the ability to retrieve resistant organoids and either characterize them via ‘omics’ approaches or test them against additional drug candidates.

Future modifications to our approach have the potential to unlock multiple additional phenotyping capabilities. For example, the 3D-trained neural network could be optimized further to track cell divisions and cell (*39*) lineages with more frequent imaging intervals (*e.g*. on the order of minutes rather than hours) for fast-dividing cell types, extending our ability to measure migration to multiple cells and enhancing resolution of cell division time measurements. In addition, we found that microwells loaded with a greater number of cells tended to grow more slowly, a phenomenon likely attributed to paracrine or juxtacrine signaling which could be investigated with the loading of cells tagged with a different fluorescent protein. For example, the addition of fluorescent tags on proteins such as MUC5AC and PGC, which are commonly used as cell type markers in mucous cells and chief cells respectively, could reveal whether certain cell types are likely to promote or suppress growth. Similarly, we could test whether adjacent apoptotic cells slow the growth of nearby cells by fluorescently labeling Annexin V (*66*). Finally, we could visualize chromosome mis-segregation errors during cell division using our H2B-mCherry nuclear reporter with higher magnification imaging (*67*).

In summary, we present a microwell-based 3D culture platform that can be easily adapted for the quantitative characterization of various image-based phenotypes. By applying this platform, we quantify cell-to-cell variability in growth rate and apical-basal polarity in two engineered gastric organoid models, and we discover potential molecular mechanisms underlying the variability in both phenotypes. Through our findings, we demonstrate the importance of single organoid measurements for characterizing morphological and molecular changes associated with disease progression, which would be impossible to accomplish through traditional bulk measurement approaches.

## Methods

### Culturing of CRISPR-engineered gastric organoids

P53KO and TP53/ARID1A DKO gastric organoids were generated as previously described (*7, 13*). Briefly, non-malignant gastric tissue from the corpus (stomach body) were obtained during sleeve gastrectomy at Stanford University Hospital under an IRB approved protocol. Wild-type gastric organoids were established followed by CRISPR/Cas9 mediated knockout of *TP53* (P53KO) (*13*).CRISPR/Cas9-mediated *ARID1A* knockout (KO) in primary *TP53*^-/-^ human gastric organoids, yielded double knockout lines (TP53/ARID1A DKO) (*7, 13*). These two organoid lines, representing pre-malignant and malignant states were used in two separate sets of experiments. In the first set (experiments #1 - #3), we conducted proof-of-principle tests to determine various types of measurements that we could perform by culturing single-cell derived gastric organoids in the microwells. Prior to the microwell experiments, organoids were cultured in 24-well tissue culture plates. These organoids were maintained in growth media containing 50% Wnt3A/R-spondin1/Noggin conditioned media produced in house, 50% Advanced DMEM/F12, 1X Penicillin/Streptomycin/Glutamine, 1X Normocin, 1X B-27 Supplement, 1X GlutaMax, 1mM N-Acetyl-L-cysteine, 500nM A83-01, 10uM SB202190, 10mM Gastrin, and 50ng/mL EGF. The media was further supplemented with 10μM Y-27632 and 2.5μM CHIR-99021 during passaging to promote stem cell survival. The culture media were refreshed every 7 days and the organoid cultures were passaged once every 12-14 days. During each passage, old media was removed and 500ul of TrypLE was added to each well to dissolve the Cultrex BME and dissociate organoids into single cells. After 30-40 minutes of incubation at 37°C, the cells were pelleted down by centrifuging at 500g for 5 minutes. The supernatant was then removed, and the cell pellet was resuspended in wash media (Advanced DMEM/F12, 1X HEPES, 1X Penicillin/Streptomycin/Glutamine) for cell counting. Cell counting was performed by combining 10ul of cell suspension with 10μl Trypan Blue, then loading the mixture into a Countess chip for automated counting using the Countess II Cell Counter (Thermo Fisher). An appropriate volume of cells was transferred to a new tube, spun down at 500g for 5 minutes, and the pelleted cells were resuspended in fresh Cultrex BME. Cells were re-plated in a new 24-well plate with 20000 cells per 40μl of Cultrex BME dome. The plate was then incubated at 37°C for 20 minutes to allow Cultrex BME to solidify; after which, 500μl fresh growth media was added. In the second set of experiments (#4 - #6), instead of using conditioned media as in the first set of experiments, we opted to use chemically defined complete growth media to prevent media batch effects from confounding our downstream analysis. The chemically defined media was composed of Advanced DMEM/F12, 1X Penicillin/Streptomycin, 1X Normocin, 1X N21-Max Supplement, 1X GlutaMax, 1mM N-Acetyl-L-cysteine, 500nM A83-01, 10uM SB202190, 10mM Gastrin, and 50ng/mL EGF. During passaging, 10μM Y-27632, 2.5μM CHIR-99021 and 200ng/ml FGF10 were added. All other culture conditions remained the same as described in the previous section.

### Lentiviral transduction to fluorescently tag organoids with histone H2B protein

Addgene plasmid #89766 expressing H2B-mCherry fusion protein was packaged into lentiviral particles (by the Gene Vector Virus Core at Stanford University). Prior to transduction, the organoids were washed with PBS and incubated with TrypLE at 37°C for 40 minutes. FBS was then added to quench the TrypLE reaction. After that, the dissociated cells were centrifuged at 500g for 5 minutes and resuspended in growth media supplemented with 10μM Y-27632. The H2B-mCherry lentiviral particles were added at an MOI of 0.1 to aliquots of 500k cells to maximize the number of successfully transduced cells harboring just one copy of the transgene. This was done to reduce insertional mutagenesis and normalize the fluorescence intensity. The cell/virus suspension was transferred to a single well of a 24-well plate, and a 1-hour spinoculation at 600g at 32°C was performed. After spinoculation, cells were incubated 37°C for four hours before being dissociated and pelleted down at 500g for 5 minutes. After which, the pellet was resuspended in Cultrex BME followed by plating onto a 24-well plate. Organoids were allowed to recover for 3-5 days until mCherry expression was visible, then the transduced cells were FACS-sorted for cells expressing positive mCherry signal. Sorted cells were allowed to recover and expand for several passages before being used for experiments.

### Microwell design and fabrication

Design files for molds containing arrays of square microwells with a variety of different dimensions (from 100 × 100 × 80μm to 1000 × 1000 × 80μm; breath × width × height) were generated in AutoCAD and printed on standard transparencies at 30000 dpi. All design files are available in Supplemental Information. Molds were created from SU-8 2100 photoresist (Microchem, Inc.) on a 4” silicon test-grade wafer (University Wafer) according to the manufacturer’s instructions. After fabrication, molds were treated with vapor deposition of 1H,1H,2H,2H-perfluorooctyl-trichlorosilane (Sigma Aldrich) under vacuum for 10 minutes. Microwell devices were made from the molds using standard soft-lithography (**Supplementary Fig. S1**). Briefly, RTV 615 precursor solutions at a ratio of 10:1 (base elastomer:curing agent) (R.S. Hughes) were mixed using a THINKY mixer with 3 minutes of mixing followed by 3 minutes of degassing, both at 2000 rpm, poured onto molds, and spun on a spin coater (Laurell Technologies) at 200 rpm with an acceleration of 133 rpm/s for 30 seconds to spread the PDMS and create a layer of PDMS approximately 0.5 mm thick (**Supplementary Fig. S2**); this thickness could be easily peeled off but remained thin enough for high-quality imaging using a long working distance objective. The PDMS was then degassed in a vacuum chamber for 15 minutes and baked at 80°C for 20 minutes. After baking, the PDMS was peeled from the mold and cut into arrays of square devices (**Supplementary Fig. S1**). Prior to use, the devices were treated with 20% oxygen plasma for 8 minutes, placed at the bottom of 12-well plates with forceps, sterilized overnight by immersion in 70% ethanol, then treated with 0.5% PBS-BSA for >1 hour to render the devices hydrophilic for cell growth.

### Cell loading into microwells

Dissociated organoid obtained during bulk culture passage were resuspended in WENR media to a concentration of 6000 cells/ml for 100μm microwells and between 600-2000 cells/ml for 200μm microwells. The cell suspension was pipetted directly onto BSA-treated microwell arrays, and plates were centrifuged at 500g for 5 min to load cells into microwells. Following cell loading, excess media was aspirated and Cultrex BME was pipetted dropwise onto the microwell arrays containing cells. Plates were incubated at 37°C to polymerize the Cultrex BME, then growth media supplemented with 10μM Y-27632 and 2.5μM CHIR-99021 and 200ng/ml FGF10 was added.

### High-resolution time-lapse image acquisition

For initial bright field imaging experiments (**Fig. 1**), organoids in microwells were grown in a standard tissue culture incubator and imaged 1X per day over 7-10 days using a Keyence BX-700 microscope. A grid acquisition with a 10X objective and 20X final magnification was performed using the built-in brightfield capability of the Keyence microscope.

High-resolution time-lapse images for experiments #1-6 (**Figs. 2-4**) were acquired on an inverted fluorescence microscope (Nikon Ti) with a motorized xy-stage (ASI MS-2000) and a camera (Andor Zyla 4.2+) set to acquire images at 2×2 binning for a final resolution of 1024×1024. Broad spectrum illumination was provided by a solid-state light source (Lumencor Sola). Images were acquired with a 10X objective at 20× final magnification. For multi-day acquisitions, organoids were kept in a stage-top incubator to maintain 37°C, 5% CO2, and 95% relative humidity. Imaging was controlled with a Jupyter notebook that used a custom Python library (AcqPak) to manage experimental acquisitions and the Micro-Manager API for hardware control; all AcqPak software is freely available for download from Github (https://github.com/FordyceLab/AcqPack). Each position in a rastered acquisition was imaged with the following filter cubes and exposures: brightfield (Semrock BRFD-A-NTE-ZERO) at 1 ms, Cy5 (Semrock 49002) at 15 ms, and mCherry at 50ms. Rasters were acquired every 2 hours.

### Initial image processing

All image processing was performed using custom Python libraries which are openly available for download from Github. To extract per-microwell information, we first stitched raw tiled images are first stitched into a single reconstructed image of the microwell array (MicrowellStitcher; https://github.com/FordyceLab/ImageStitcher). The pixel locations of the array corners are used to rotate the image so the array edges are parallel with the image edges. Subarrays containing either 400 microwells (100um microwells) or 100 microwells (200um microwells) are extracted from the stitched array image using the corner pixel locations (MicrowellProcessor; https://github.com/FordyceLab/MicrowellProcessor), and image prediction for nuclear segmentation is performed with a trained deep learning model on the subarrays (Wellception; https://github.com/juliaschaepe/wellception).

### Deep learning model training and testing

To enable automated quantification of organoid growth from microwell images, we first trained a deep learning model (*38, 39*) to predict fluorescent organoid nuclei. To construct the dataset used to train the model, individual microwell images of 114×114 pixels from experiment #1 were saved as .npz files, and each image was randomly assigned to either training data (80%), validation data (20%). Additional images from experiments #1 - #3 were used as test data (306, 348, 258 images respectively). We used automated quality control to remove microwell images containing >8 cells after initial testing revealed images with higher cell numbers were challenging for annotators to label manually (data not shown), leaving a final dataset of 2711 training images, 542 validation images and 912 test images. The Caliban desktop module (*39*) was used to manually label individual nuclei in each image, and the annotations were saved to the .npz files. We then used deepcell-tf (https://github.com/vanvalenlab/deepcell-tf) to train a deep learning model, with model weights stored in a hdf5 file, and performance was measured on the validation and test sets using the deepcell-toolbox (https://github.com/vanvalenlab/deepcell-toolbox)(*38, 39*).

To predict nuclei in microwell images, the organoid segmenter was initialized with trained model weights stored in an hdf5 file. The segmenter took in input images and preprocessed them with histogram normalization using a kernel size of 32 × 32. Each macrowell was imaged in 9-15 subarrays, with each 2280 × 2280 pixel subarray containing 400 microwells of 114×114 pixels each (100μm microwells) or 100 microwells of 224 × 224 pixels each (200μm microwells). Each subsection was first rescaled by a factor of two before being passed into the model. Model outputs were postprocessed with a watershed filter with a detection threshold of 0.25, a distance threshold of 0.1 and a minimum distance of 2.5. The model was run on a NVIDIA GPU. Nuclear predictions were saved as an hdf5 file containing all subarrays coming from a single microwell array. A .csv file containing summary statistics of the predictions for each timepoint and microwell (indexed by pixel location), including the predicted number of nuclei, area of each nucleus, the total area of the organoid, and centroid locations for each organoid, alongside pertinent metadata such as timepoint, microwell ID, mutant, and microwell pixel location was generated.

### Quantification of organoid growth rates using deep learning

To quantify organoid growth rates, we first used the extracted deep learning predictions to identify microwells initially loaded with a ≥1 cells for which the model identified ≥n+1 cells at the final timepoint in the timecourse. Total predicted organoid nuclear area was plotted as a function of time for up to 60 timepoints to include only timepoints with >8 cells. A model of exponential growth was then fit to the data, and parameters corresponding to initial area and growth rate as the percent change in area per day were extracted from the exponential model. To compare growth rate distributions across mutants and experiment, a one-way ANOVA with three variables was performed, and Bonferroni correction was used to account for multiple hypothesis testing.

### Polarity measurements with confocal imaging

Initial confocal imaging to assess apical-basal polarity was performed in microwells. Growth media was aspirated from plates, arrays were washed with 1X phosphate-buffered saline (PBS), the arrays were incubated in BD Cytofix/Cytoperm Fixation/Permeabilization solution for 30min at 4°C, then cells were washed 3× with 1X BD Perm/Wash buffer, with 10 minute incubations at room temperature between washes. Fixed cells were then stained with DAPI at 300 nM concentration diluted in PBS solution and Alexa Fluor 647 phalloidin at 165 nM concentration diluted in PBS solution, then incubated at room temperature in the dark for 1 hr. Cells were washed with PBS solution before imaging. Imaging was performed with a Leica SP5 upright multi-photon confocal microscope with a HCX APO L20×/1.00 water immersion objective. Organoid images were acquired in a z-stack with images taken every 10 μm. Representative single-plane images were extracted using Volocity software.

### Polarity measurements with live-cell fluorescence imaging

For experiments #1 - #3, 1X SiR-actin dye (Cytoskeleton Inc.) and 1X verapamil were added to growth media before addition to macrowells at the start of the timecourse. For experiments #4 - #6, growth media was aspirated from microwells at the final timepoint of the timecourse, and growth media supplemented with 1X SiR-actin dye and 1X verapamil was added to macrowells. This was done in order to reduce the extrinsic effects introduced by the addition of the actin dye on the growth measurements of the organoids. Imaging, image processing, and individual microwell extraction were performed as described above. Blinded manual polarity classification was performed on a subset of images from the final timepoint of experiment #1, with 408 microwells classified for TP53KO and 403 microwells for DKO. Organoids were classified as “normal”, “abnormal”, or “unknown”. To be classified as “abnormal”, organoids were required to display 2 of the following 3 characteristics: (1) disorganized actin signal not restricted to organoid lumens, (2) lack of a central lumen ringed with actin, and/or (3) the presence of multiple small, disorganized lumens. Aspiration of organoids with varying polarity was performed at the end of experiment #6, using the same classification requirements.

### Shallow WGS of bulk organoids

The TP53KO and DKO organoids were harvested using TrypLE solution in a fashion similar to organoid passaging, prior to the start of experiments #4 and #6. The cells were then lysed and the nucleic acid was extracted using Qiagen Allprep DNA/RNA Mini Kit (Qiagen). Aliquots of DNA were sent to Novogene Co. for the construction of sequencing library and shallow WGS at 1X coverage. Sequencing reads were aligned to the hg38 human reference genome using BWA (*68*). Samtools (*69*) was used to convert the alignment files into bam format with indexes. The bam files were subsequently analyzed with QDNAseq (*70*) to infer copy-number variations using 50kb read bin size and median normalization. The output was log2 transformed and visualized using a custom R script.

### Single organoid retrieval from microwell

A syringe pump (Harvard Apparatus Pump 11 Elite) was used to drive a 1 ml syringe fitted with a blunt Luer probe tip. The syringe was attached to a 255μm ID/510μm OD PEEK tubing via a short 1 cm Tygon splint. The PEEK tubing was then threaded through a tightly-fitting 20μl pipette tip such that approximately 0.5cm extended past the tip. The splint and tip connections were secured with super glue. The tip was affixed to the microscope condenser z-stage using a 3D-printed liquid light guide holder (**Supplementary Fig. S8**). The syringe and tubing were primed with phosphate-buffered saline (PBS). The touch-down position of the tip was adjusted to the center of a live field of view under 4X magnification using thumbscrews on the condenser slot; this position was noted by drawing a reference box around the tip location in the Micro-Manager GUI. To pick an organoid of interest, the xy-stage was moved until the organoid was centered in the reference box. The tip was then lowered to the microwell surface using the condenser z-stage. The syringe pump was then used to withdraw the organoid into the tip; if necessary, adherent organoids were loosened by toggling withdraw/infuse on the syringe pump and/or by incubation with TrypLE for 5 minutes. The tip containing the aspirated organoid was then positioned just above an empty container (e.g. a well of a multiwell plate) and dispensed using the infuse button on the syringe pump into a 1.5ml centrifuge tube. A total of 20 organoids were retrieved from the DKO culture from experiment #6 in microwells. Ten organoids had normal apical basal polarity as evidenced by the actin ring with a large lumen (**Fig. 4B**). Ten organoids had abnormal apical basal polarity as defined throughout based on the presence of two of the three criteria: (1) disorganized actin signal not restricted to organoid lumens, (2) lack of a central lumen ringed with actin, and/or (3) the presence of multiple small, disorganized lumens.

### Dual ATAC/mRNA library preparations and sequencing of single retrieved organoids

Each single organoid retrieved from microwell was subjected to dual ATAC/mRNA sequencing library preparation following a recently published protocol that was designed for low input (*57*). Briefly, the organoid was spun down at 500g for 5 minutes using a benchtop centrifuge. After removing supernatant, the organoid was then resuspended in a direct permeabilization/tagmentation mastermix (25μl TD buffer (from Illumina Nextera XT DNA Library Prep Kit), 2.5μl Tn5, 16.5μl DPBS, 0.5μl 1% digitonin, 0.5μl 10% Tween-20, 2.5μl RNase inhibitor and 2.5μl nuclease-free water) without prior nuclei isolation. The reaction mixture was incubated in a 37°C water bath for 30 minutes with occasional hand-pipetting. At the end of incubation, 2.5μl of stop buffer containing 10mM EDTA and 0.5M lithium chloride was added to neutralize the permeabilization/tagmentation reaction. The tagmented cells were then lysed using 100μl of Lysis/Binding Buffer from the Dynabeads mRNA Direct Micro Kit (Invitrogen). After lysis, 20μl of pre-washed Dynabeads Oligo-dT beads were added to the lysate which was thereupon incubated at room temperature for 5 minutes to allow mRNA to anneal to the Oligo-dT. The beads with annealed mRNAs were separated from the supernatant using a magnetic rack. The supernatant which contained genomic DNA was transferred to a new tube for subsequent genomic DNA extraction using Qiagen MinElute PCR Purification Kit. Meanwhile, the beads-mRNA complex was resuspended in 20μl of reverse transcription mastermix (without any primer) from Superscript IV First Strand cDNA Synthesis Kit (Invitrogen). The Oligo-dT on beads served as primers for the reverse transcription reaction. The reaction mix was incubated at 50°C for 5 minutes then at 55°C for an additional 10 minutes. The resulting cDNA/mRNA hybrid was thus covalently bound to the magnetic beads. The beads were washed twice with 100μl ice-cold 10mM Tris-HCI (pH 7) and resuspended in 5μl of Tris-HCl. Sequencing library preparation of the cDNA was performed on beads using Nextera XT DNA Library Prep Kit where the reagent volumes were halved (keeping the reagent ratio consistent). Meanwhile, the previously tagmented ATAC-DNA was amplified using Q5 High-Fidelity 2X Mastermix (NEB) with universal i5 and indexed i7 adapters. PCR was performed with 18 cycles due to the low input nature of the samples. Both ATAC and cDNA libraries were cleaned using AMPure XP beads according to manufacturer’s recommendations. The libraries were then sent to Novogene Co. for sequencing on the Illumina Novaseq platform with 75bp paired-end reads.

### Bioinformatic analysis of ATAC-sequencing libraries

The raw sequencing reads were first processed with Trim-Galore to remove Illumina adapter sequences. The Nextflow atacseq pipeline (nf-core/atacseq) was used to process the sequencing trimmed reads with default parameters. Briefly, reads were aligned to hg38 human reference genome using BWA. The alignment files were further processed with Picard and Samtools to mark duplicate reads and to create bigWig files for downstream analysis and data visualization. Open chromatin peaks were called using MACS2 and annotated with HOMER. Due to low sample input, the number of PCR cycles used to generate the library was more than the recommended number, and this resulted in higher read duplication rate. As a result, the data was subjected to additional filters: 1) samples with less than 60000 deduplicated, uniquely mapped reads, and 2) samples with less than 500 total peaks were removed. The final dataset consisted of seven organoids each with normal and abnormal apical-basal polarity (**Fig. 5C**). The peak regions were then analyzed in DESeq2 to determine differential accessibility regions (DARs) (*71*). The transcription factor footprinting analysis was performed using TOBIAS with default parameters (*72*). The DARs were subjected to gene ontology (GO) analysis using default parameters (*73–75*). The GO terms with a p-value <0.01 were retained for Revigo analysis to remove redundant GO terms (*76*).

### Bioinformatic analysis of mRNA-sequencing libraries

The raw sequencing reads were first processed with Trim-Galore (*77*) to remove Illumina adapter sequences. The trimmed reads were then aligned to hg38 human reference transcriptome using STAR (*78*), and processed using RSEM to calculate gene expressions in each sample (*79*). The data was further examined with PCA analysis to remove outlier samples. In addition, the genes with zero readcount in more than or equal to 30% of the samples were removed from the differential analysis to ensure that the analysis was not skewed by missing data. DESeq2 was used to determine differential gene expression between the normal and abnormal groups. Gene ontology analysis was performed in similar manner as described for ATAC-sequencing libraries with one modification: GO terms with a p-value <0.05 were retained for Revigo analysis due to a low number of differentially expressed genes (**Supplementary Fig. S9**).

## Data Availability

Raw sequencing data for shallow WGS and dual ATAC/RNA sequencing is available on SRA with accession number PRJNA858865 at the NCBI Short Read Archive (https://www.ncbi.nlm.nih.gov/bioproject/PRJNA858865/). Other data, including image files for experiments #1-6, labeled training, validation, and test set data for deep learning model training, processed data files, and per-experiment and per-macrowell summary reports are available through OSF at https://osf.io/r4s3m/.

## Acknowledgments

The authors thank the Cell Sciences Imaging Facility and Genetics Bioinformatics Service Center at Stanford University for assistance in confocal microscopy and computational resources respectively. This work was supported by the NIH Director’s Pioneer Award (DP1CA238296) to C.C. and the NIH Director’s New Innovator Award (DP2GM123641) to P.M.F. C.C and P.M.F are Chan Zuckerberg Biohub Investigators. We thank members of the Curtis and Fordyce lab for helpful feedback on this manuscript.

## Author Contributions

A.S., W.W, K.K., C.C., and P.M.F. conceptualized the initial research idea. A.S. and W.W. performed the microwell experiments. A.S. tested and made the microwell devices and performed image processing bioinformatic analysis. W.W. performed organoid picking experiments, made single-organoid sequencing libraries and performed bioinformatic analysis. S.L. and D.M. set up the microscope, performed initial testing, and implemented the organoid-picking platform. T.V. performed manual labeling of deep learning training and test data. V.C. performed initial growth and cell seeding experiments. D.V.V., J.S. and A.S. set up the deep learning pipeline. K.K. and Y.L. performed the organoid gene-editing. A.S., W.W., C.C., and P.M.F wrote the manuscript.

