## Supplemental Information for "A microwell platform for high-throughput longitudinal phenotyping and selective retrieval of organoids"

**This Supplementary Material file includes:**

Supplementary Figs. S1 to S9

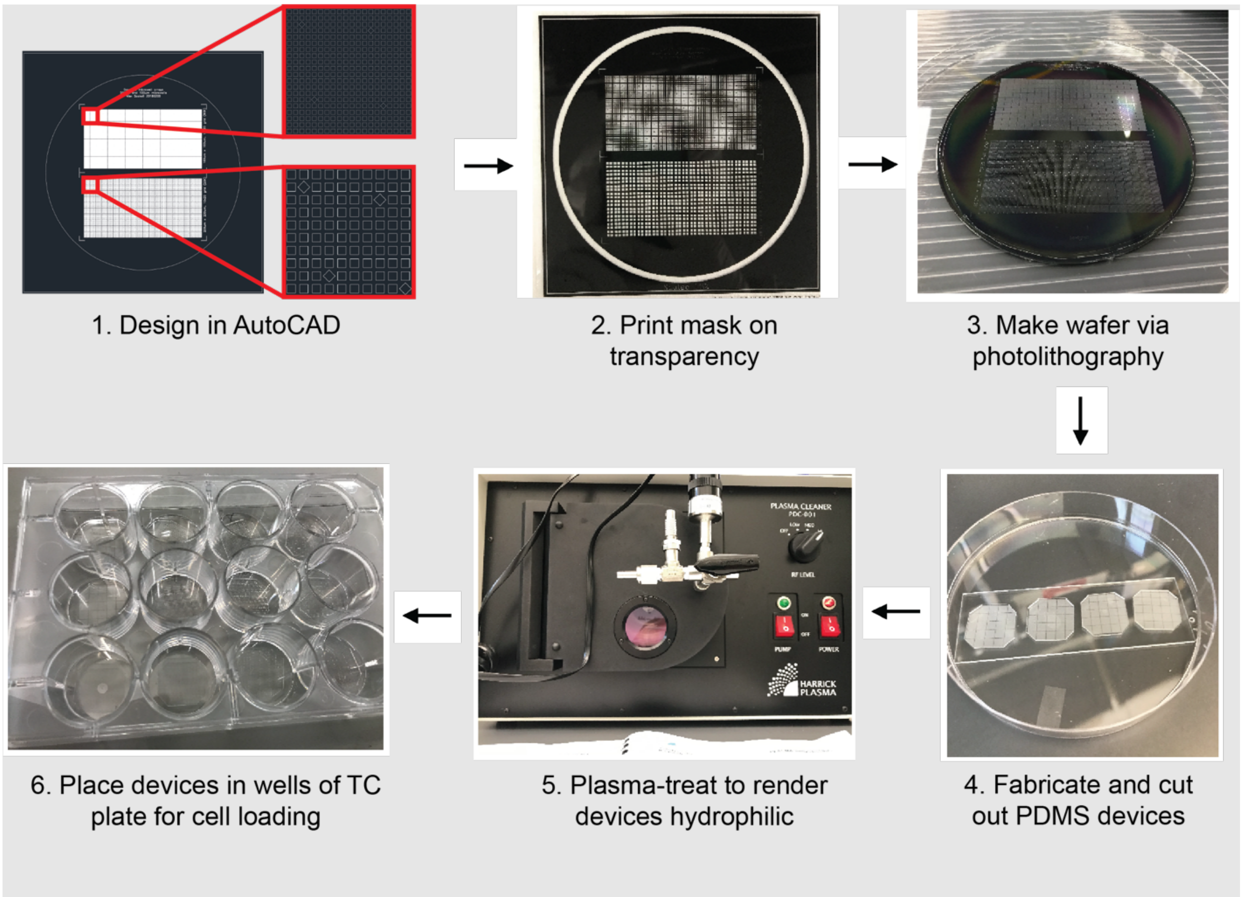


**Supplementary Figure S1. PDMS microwell array fabrication.** Microwell arrays of pre-defined dimensions (100 x 100 x 80 or 200 x 200 x 80 µm) were designed in AutoCAD and printed on transparencies that were used to fabricate silicon wafer molding masters via standard photolithography. Microwell devices were created by spin-coating PDMS onto molding masters and cutting out individual microwell arrays. After fabrication, devices were exposed to oxygen plasma to render surfaces hydrophilic and then inserted into the bottom of individual 'macrowells' within standard 12 well tissue culture plates.

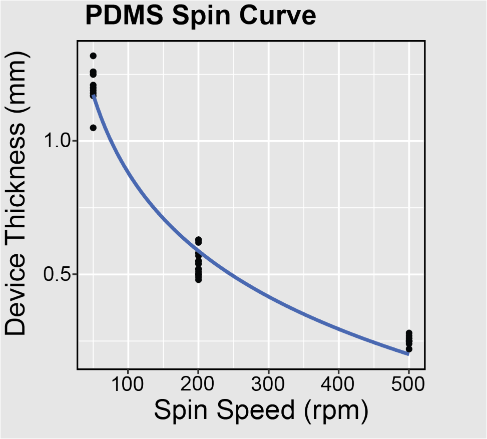


**Supplementary Figure S2. PDMS spin curve.** Relationship between final device thickness and PDMS spin speed. Spin coating at 200 rpm yielded 0.5 mm thick devices used in downstream experiments.

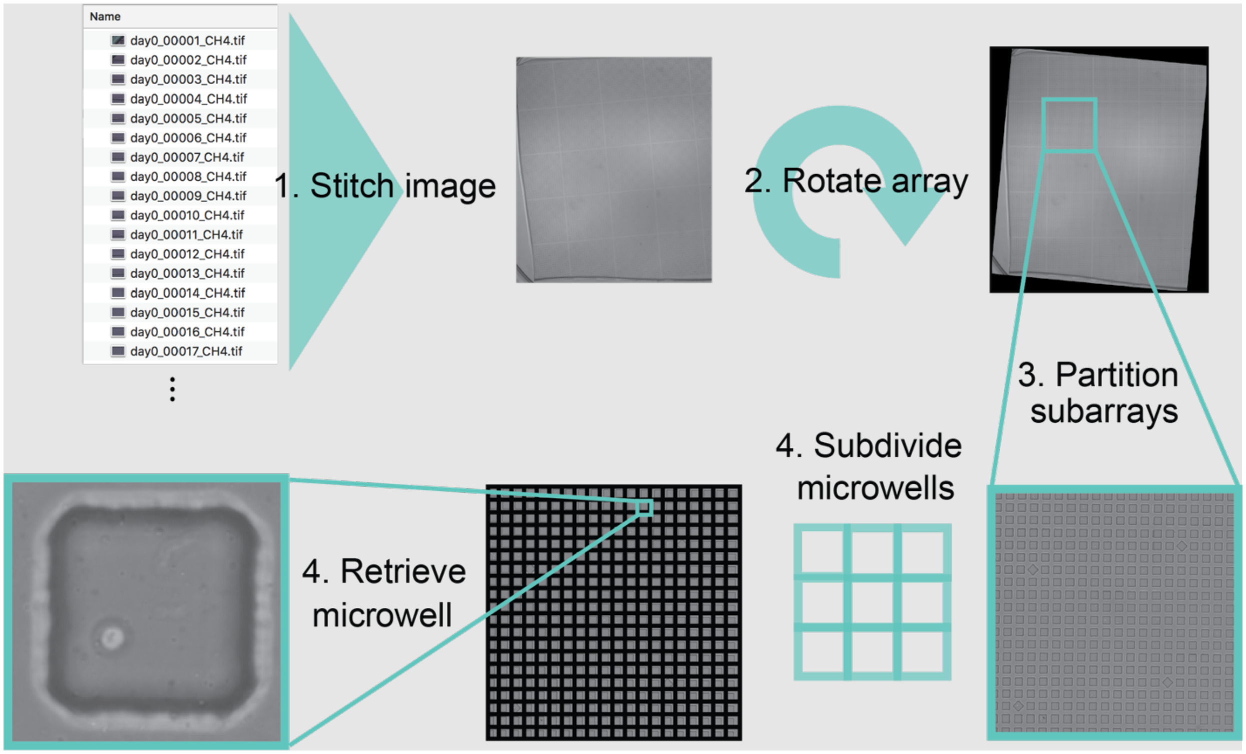


**Supplementary Figure S3. Image analysis pipeline.** (1) Devices are imaged via a tiled acquisition at each timepoint and then stitched to form a single image of the entire device at each timepoint, (2) stitched images are rotated to align device microwells perpendicular to the borders of each image, (3) rotated images are subdivided by subarrays, (4) subarray images are further subdivided by microwells, then individual microwell images are extracted from the subarrays, and images over time are collated for each microwell . Processing is done for all timepoints in parallel and downstream image processing allows identification of microwells containing cells.

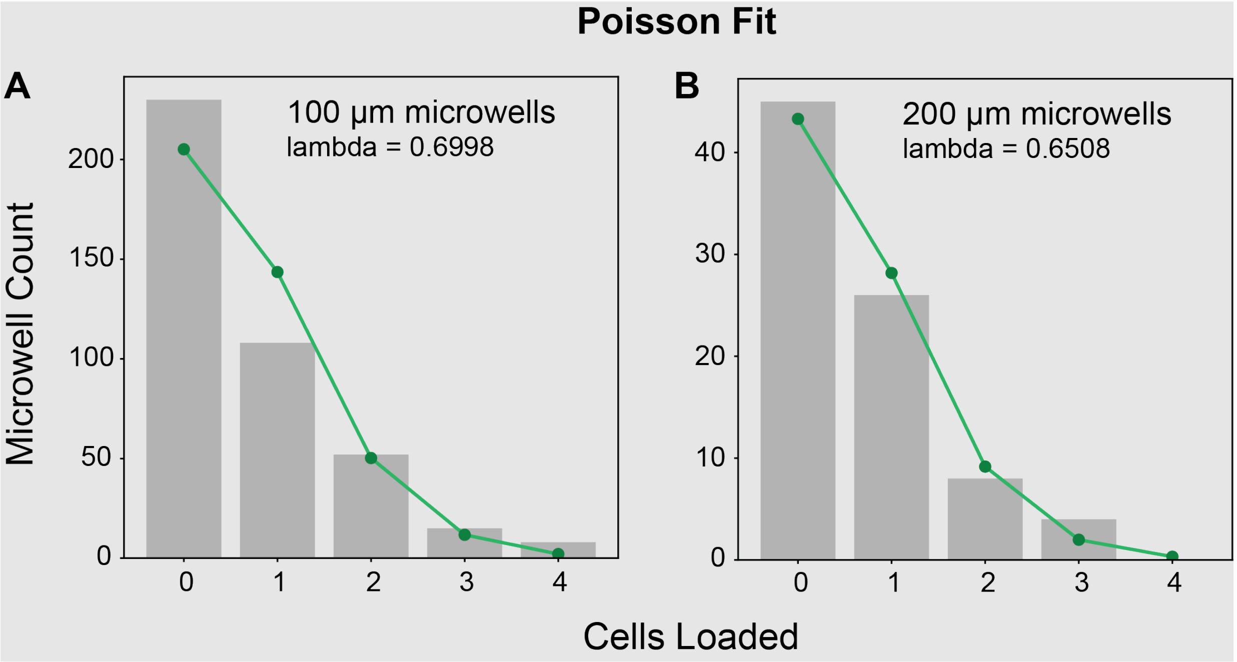


**Supplementary Figure S4. Cell loading statistics.** Number of microwells containing 0, 1, 2, 3, or 4 cells post-loading for 100 µm x 100 µm **(A)** and 200 µm x 200 µm **(B)** microwell arrays; green line and markers indicates best fit Poisson with annotated fit parameter.

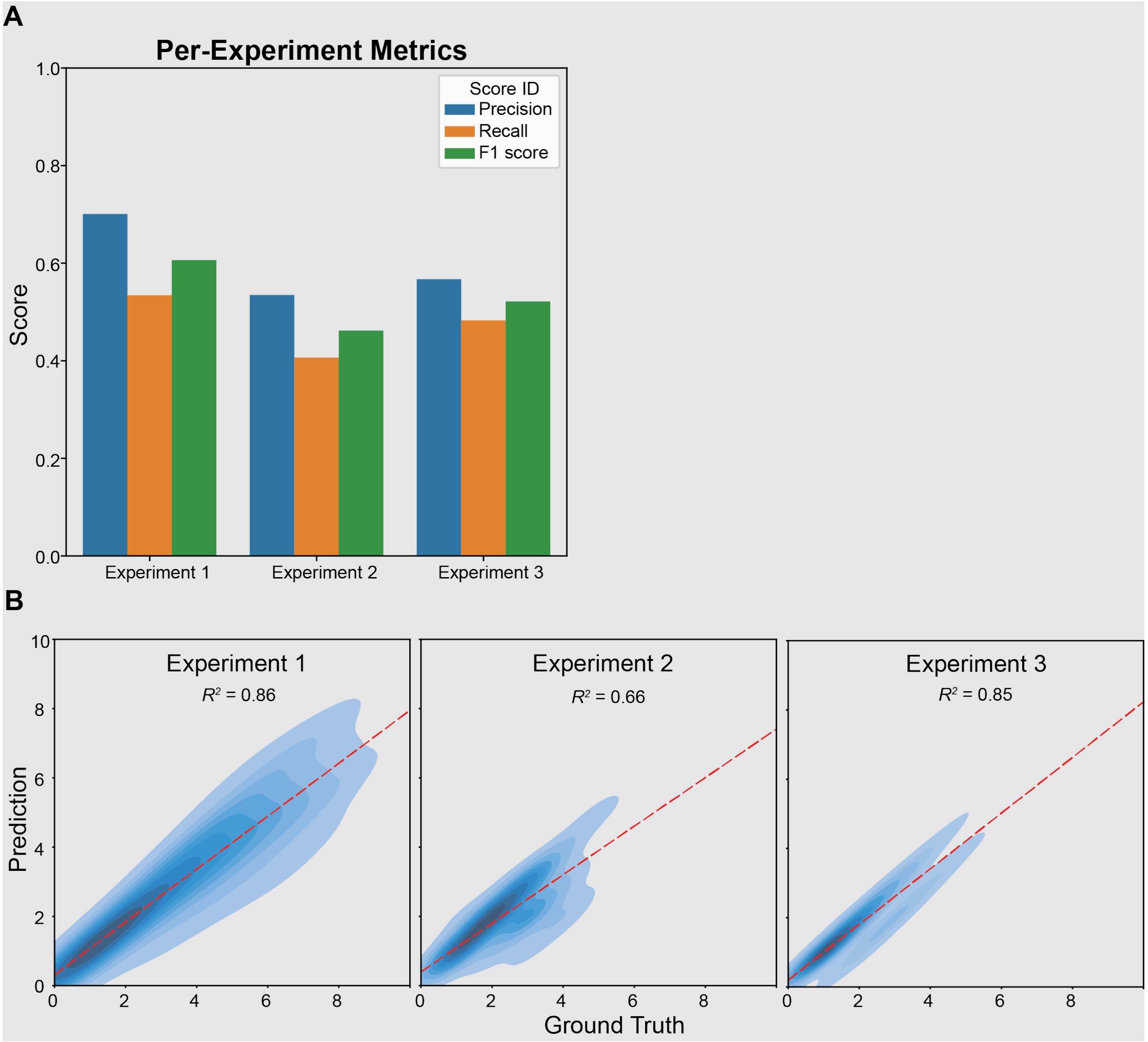


**Supplementary Figure S5. DeepCell training and testing. (A)** Per-experiment precision, recall, and F1 scores for DeepCell performance across the Experiments #1-3 test set; scores were highest for Experiment 1, as expected given that data from this experiment were used to train the model. **(B)** Correlations between per-microwell DeepCell-predicted cell counts *vs.* manually labeled 'ground truth' cell counts across Experiments #1-3; dashed red line indicates linear regression with annotated Pearson correlation coefficients.

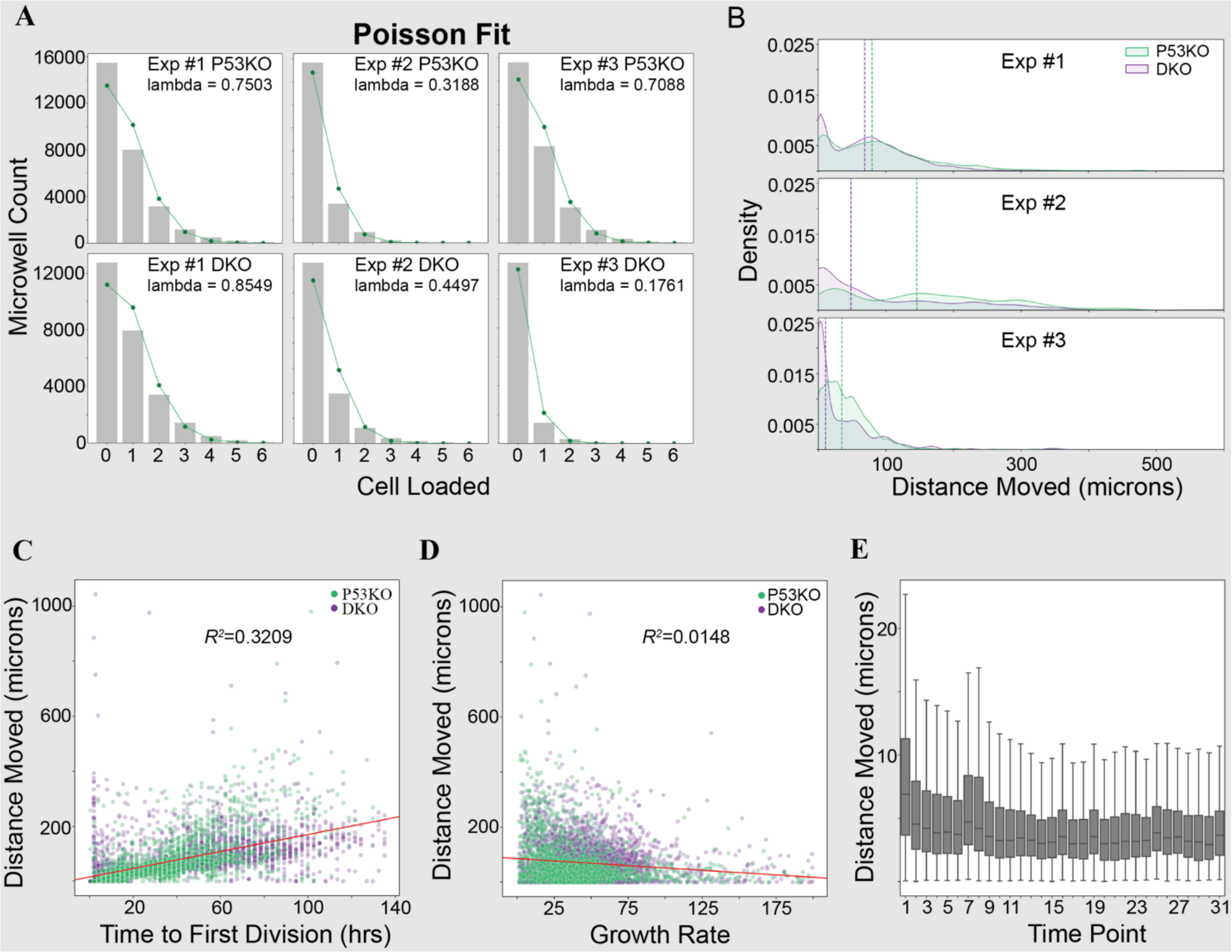


**Supplementary Fig. S6. Additional DeepCell-enabled phenotypic characterization of organoids grown in experiments #1 - #3. (A)** Number of microwells containing 0-6 cells after loading for ΔP53 (top) and DKO (bottom) cell lines; green line and markers indicate best fit Poisson with indicated fit parameter. **(B)** Density plots showing distance moved by single cells prior to first division for ΔP53 (green) and DKO (purple) cell lines; dashed lines indicate population median. **(C)** Distance moved *vs.* time to first division for single ΔP53 (green) and DKO (purple) cells loaded within microwells; red line indicates linear regression with annotated Pearson correlation coefficient. **(D)** Distance moved *vs.* calculated growth rate for single ΔP53 (green) and DKO (purple) cells loaded within microwells; red line indicates linear regression with annotated Pearson correlation coefficient. **(E)** Boxplots showing distribution of distances moved by all single cells at each time point prior to the first cell division.

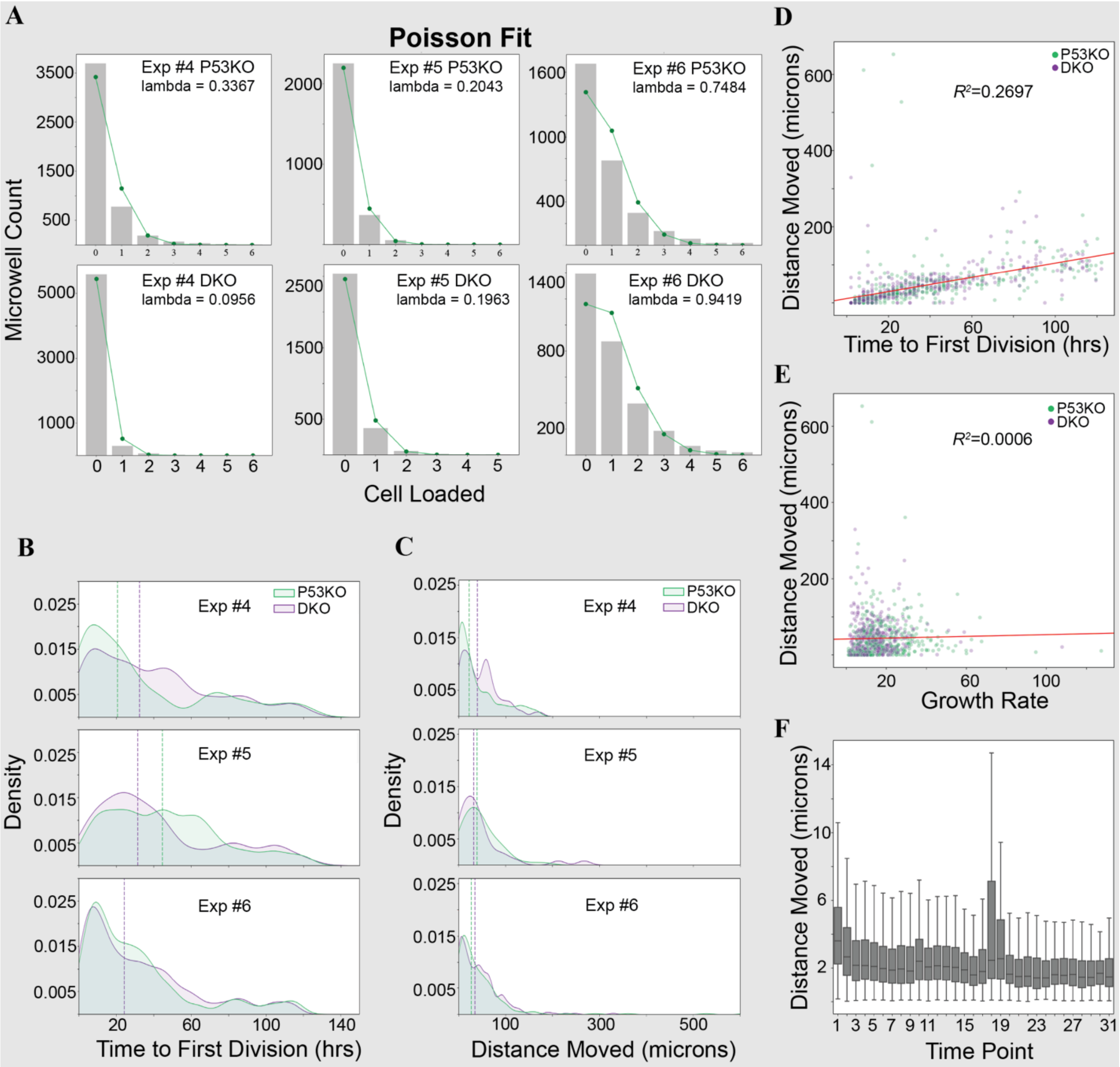


**Supplementary Figure S7. Additional DeepCell-enabled phenotypic measurements of organoids grown in experiments #4 - #6. (A)** Number of microwells containing 0-6 cells after loading for ΔP53 (top) and DKO (bottom) cell lines; green line and markers indicate best fit Poisson with indicated fit parameter. **(B)** Density plots showing time to first division ΔP53 (green) and DKO (purple) cell lines; dashed lines indicate population median. **(C)** Density plots showing distance moved by single cells prior to first division for ΔP53 (green) and DKO (purple) cell lines; dashed lines indicate population median. **(D)** Distance moved *vs.* time to first division for single ΔP53 (green) and DKO (purple) cells loaded within microwells; red line indicates linear regression with annotated Pearson correlation coefficient. **(E)** Distance moved *vs.* calculated growth rate for single ΔP53 (green) and DKO (purple) cells loaded within microwells; red line indicates linear regression with annotated Pearson correlation coefficient. **(F)** Boxplots showing distribution of distances moved by all single cells at each time point prior to the first cell division.

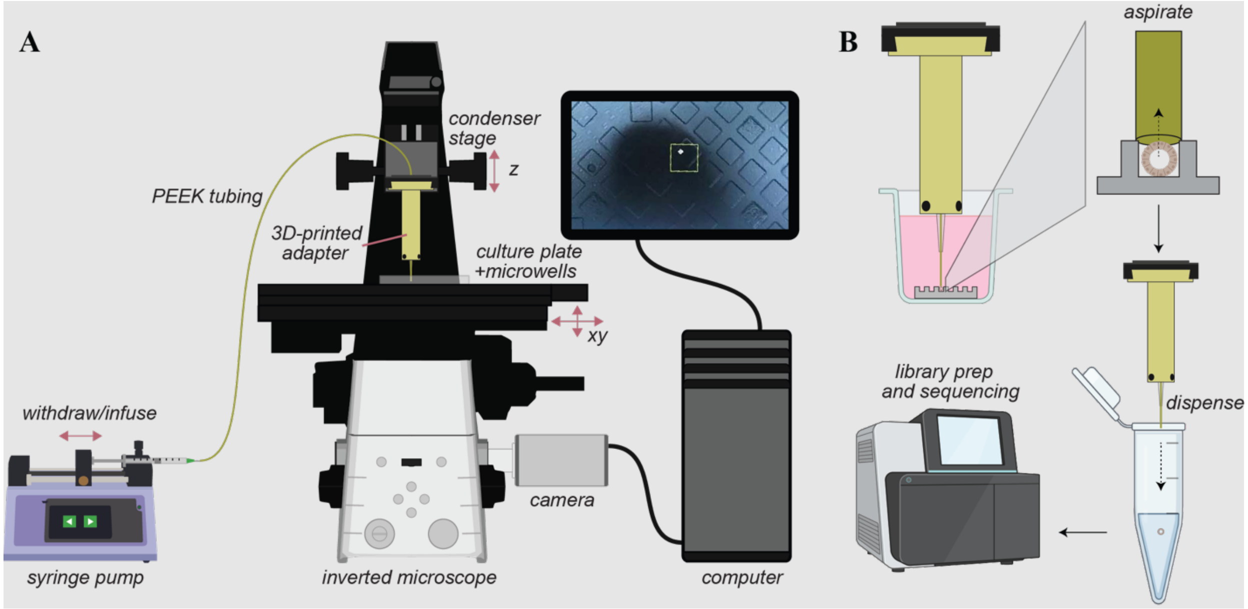


**Supplementary Figure S8. Schematic of single-organoid retrieval for sequencing. (A)** Schematic showing inverted epifluroescence microscope with 3D printed adapter and syringe pump for organoid retrieval. **(B)** Schematic showing process for retrieving organoids of interest from microwells for sequencing.

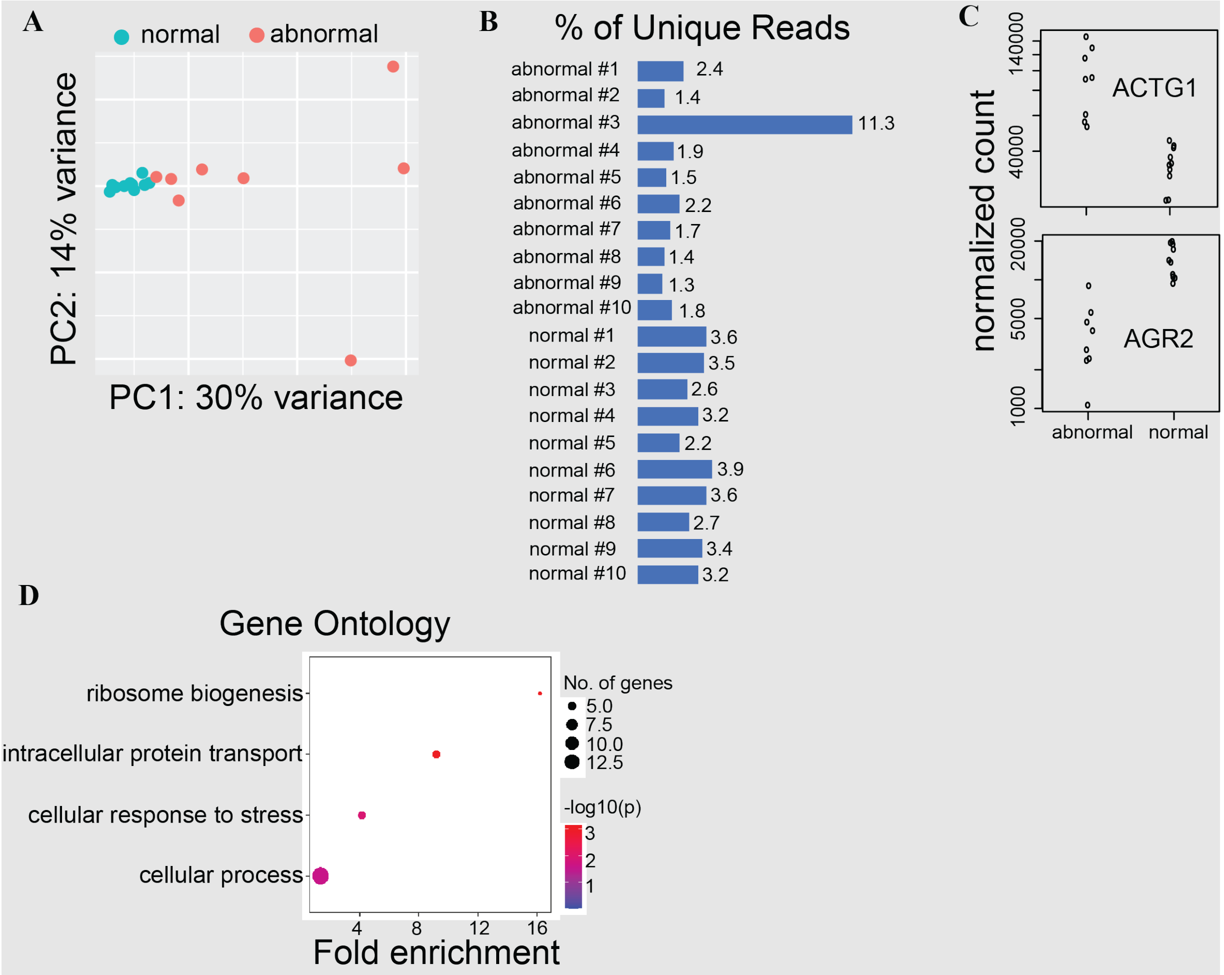


**Supplementary Figure S9. Analysis of single-organoid RNAseq. (A)** Principal component analysis of RNAseq data indicating positions of normal (green) and abnormal (red) organoids along with the percentage of variance explained by the first two principal components. **(B)** Percentage of unique sequencing reads for each organoid. **(C)** Normalized read counts for ACTG1 (top) and AGR2 (bottom) transcripts for abnormal and normal cells. **(D)** Fold-enrichment of differentially-expressed gene as a function of gene ontology term demonstrating enrichment of genes in intracellular protein transport and stress response, associated with actin-mediated functions.
